# The unfolded protein response in alveolar type II cells is a therapeutic target in ischemia-reperfusion injury and chronic lung allograft rejection

**DOI:** 10.64898/2026.11.25.769302

**Authors:** Daniel Vosoughi, Amanda Luca, Tanroop Aujla, Amit Patel, Allen Duong, Ke Fan Bei, Mohamad Elbayomi, Nicole Chrysler, Andrzej Chruscinski, Mingyao Liu, Shaf Keshavjee, Marcelo Cypel, Tereza Martinu, Stephen Juvet

## Abstract

Although lung transplantation is life saving for patients with end-stage lung disease, its long-term outcome is poor owing to an inexorable process of chronic rejection that culminates in chronic lung allograft dysfunction (CLAD) and death or the need for re-transplantation at a median of 6 years post-transplant. Current immunosuppression fails to prevent CLAD, suggesting the existence of alternative pathways not susceptible to these drugs. Lung-directed autoantibodies have been shown to predict CLAD and reduced post-transplant survival. Here, we observed that serum and bronchoalveolar lavage antibodies to GRP78, a master regulator of the unfolded protein response, identified patients at increased risk for CLAD and death. Anti-GRP78 antibodies were associated with upregulation of *HSPA5*, the gene encoding GRP78, in alveolar type II cells from human lung allografts, and with translocation of GRP78 to the cell surface, a hallmark of the unfolded protein response. Cell surface GRP78 expression was triggered by lung ischemia-reperfusion injury and was also prominent during the development of ischemia-reperfusion injury-augmented chronic rejection in a mouse lung transplant model. Treatment with the chemical chaperones 4-PBA and TUDCA, which are known to ameliorate endoplasmic reticulum stress during the unfolded protein response, reduced both ischemia-reperfusion injury and chronic allograft rejection *in vitro* and in mice. Taken together, our data implicate alveolar type II cell stress in the pathogenesis of lung ischemia-reperfusion injury and fibrotic chronic lung allograft rejection and suggest that this phenomenon can be targeted for clinical benefit.

**One Sentence Summary:** Circulating and intragraft antibodies reveal an unfolded protein response in alveolar type II cells during ischemia-reperfusion injury and chronic lung allograft rejection, which can be modulated therapeutically using chemical chaperones.

## INTRODUCTION

Despite improvements in short-term survival following lung-transplantation (LT), chronic lung allograft dysfunction (CLAD) remains the principal limitation to long-term survival (*1, 2*). The consequent median survival rate of only 6 years places LT survival as the worst among solid organ transplants (*1*). Although attributed to numerous risk factors, CLAD pathogenesis remains poorly understood. Its two major phenotypes are bronchiolitis obliterans syndroms (BOS) – obstructive physiological decline and small-airway fibrosis – and RAS – restrictive decline, diffuse alveolar injury, parenchymal fibrosis, and worse prognosis (*3*). RAS has been compared with other fibrotic lung diseases, including idiopathic pulmonary fibrosis (IPF), although with distinct histology – usual interstitial pneumonia in IPF (*4*) and alveolar fibroelastosis in RAS (*3*). Both RAS and IPF share a restrictive physiology, and numerous cellular and molecular features (*5, 6*), suggesting shared pathogenic mechanisms.

Alloimmune recognition of donor human leukocyte antigen (HLA) is an established risk factor for CLAD (*7*). Its humoral arm, characterized by donor HLA-specific antibodies (DSAs), confer a significantly increased risk of CLAD and are widely regarded as the principal humoral challenge following LT (*7–9*). Yet, autoantibodies may arise against non-HLA targets, and their contribution to CLAD has been relatively underappreciated. The mechanisms by which self-tolerance breaks down leading to humoral autoimmunity remain incompletely understood. In a murine model, central deletion of T cells specific for the bacterial Cre antigen, used as a lung-restricted model antigen, was inadequate, and the peripheral tolerance that remained was readily disrupted by successive immune challenges (*10*). Tolerance to lung antigens may therefore be inherently precarious after transplantation, when it may be susceptible to disruption by allograft inflammation. The exposure of antigens normally sequestered from immune surveillance participates in the ensuing autoimmune response. The two self-antigens whose antibodies have been best associated with subsequent CLAD fit this description: collagen V (ColV), a minor collagen buried within type I collagen fibrils, and the intracellular cytoskeletal protein K-α1-tubulin (KAT), both normally sequestered from the immune system (*11–13*). Autoimmunity to both targets has been linked to ischemia-reperfusion injury (IRI) (*14–17*), an early allograft insult characterized by a sterile inflammatory process (*18, 19*). ColV and KAT need not be unique in this respect; IRI might comparably expose other sequestered antigens whose recognition begins at engraftment and persists toward CLAD, and examining such targets could offer a window onto the antigens and pathways involved in the syndrome.

In this study, we applied our microarray-based high-content antibody screen (*20*) to patient serum and bronchoalveolar lavage (BAL) to identify glucose-regulated protein 78 (GRP78) as a major target. Anti-GRP78 IgG were consistently elevated in patients who later developed CLAD, and those with IPF prior to transplant. Cell-surface GRP78 (csGRP78) exposure has been linked to the presence of anti-GRP78 IgG across numerous pathological contexts (*21–30*). In CLAD and IPF, csGRP78 was elevated in alveolar type II (AT2) cells relative to donor lung tissue, implicating this finding as a shared feature of fibrotic lung disease before and after transplantation.

We therefore hypothesized that AT2 dysfunction may begin as early as the IRI sustained at implantation. To determine whether this response could be targeted, we used 4-PBA and TUDCA – chemical chaperones known to stabilise protein folding, dampen ER-stress signalling, and reduce collagen deposition in bleomycin-induced lung fibrosis (*31*). In a murine hilar-clamp model of unilateral IRI, intravenous dosing before clamping and again after clip release reduced lung wet-to-dry ratio and epithelial csGRP78.

Epithelial csGRP78 positivity tracked with the severity of fibrotic injury in a murine orthotopic left LT (OLT) model. Sustained post-transplant 4-PBA-TUDCA treatment reduced epithelial csGRP78, with downward trends in rejection, interstitial inflammation, and vascular fibrosis. Together, these data position AT2 ER stress and GRP78 desequestration as early reperfusion-associated events that track with later CLAD pathology and IPF. They nominate csGRP78 and the associated anti-GRP78 IgG response as candidate markers of perioperatively initiated AT2 dysfunction and suggest ER-stress mitigation as a potential intervention.

## RESULTS

### Anti-GRP78 IgG were significantly elevated prior to the onset of CLAD and in patients with interstitial lung disease

Using a novel method for the multiplexed detection of IgM, IgG, and IgA antibodies (Figure 1A) against a custom panel of antigens enriched for those localized to the lung (Supplemental Table 1), we interrogated autoantibodies in serum collected prior to LT paired to serum and BAL collected approximately 6-months post-LT (Feature Discovery Cohort [LT-FDC] cohort, Supplemental Figure 1A). Between-group analysis of average serum and BAL antibody levels across all isotypes at each timepoint revealed that these were similar between patients who developed CLAD by five years and those who remained CLAD-free (Figure 1B). However, Significance Analysis of Microarrays (SAM) analysis revealed numerous specific antibodies elevated in patients with subsequent CLAD including six in both serum and BAL (Figure 1C). We previously demonstrated that individual serum autoantibody levels were only modestly associated with future CLAD, while their integration into a combinatorial signature dramatically improved discrimination (*32*). Using our previously described autoantibody scoring method (*32*) in conjunction with cross-validated greedy backward stepwise selection limited to antibodies elevated in both serum and BAL, we demonstrated the utility of these features as a biomarker of future CLAD (Figure 1D). The smallest set of features that yielded the highest cross-validated area under the receiver operator characteristic curve (AUROC) were BAL anti-Collagen IV IgG, BAL anti-GRP78 IgG, and serum anti-club cell secretory protein (CCSP) (AUROC = 0.70). Of these, backward selection resulted in BAL anti-GRP78 IgG as the final feature that held substantial discriminative power (AUROC = 0.63).

**Figure 1.**
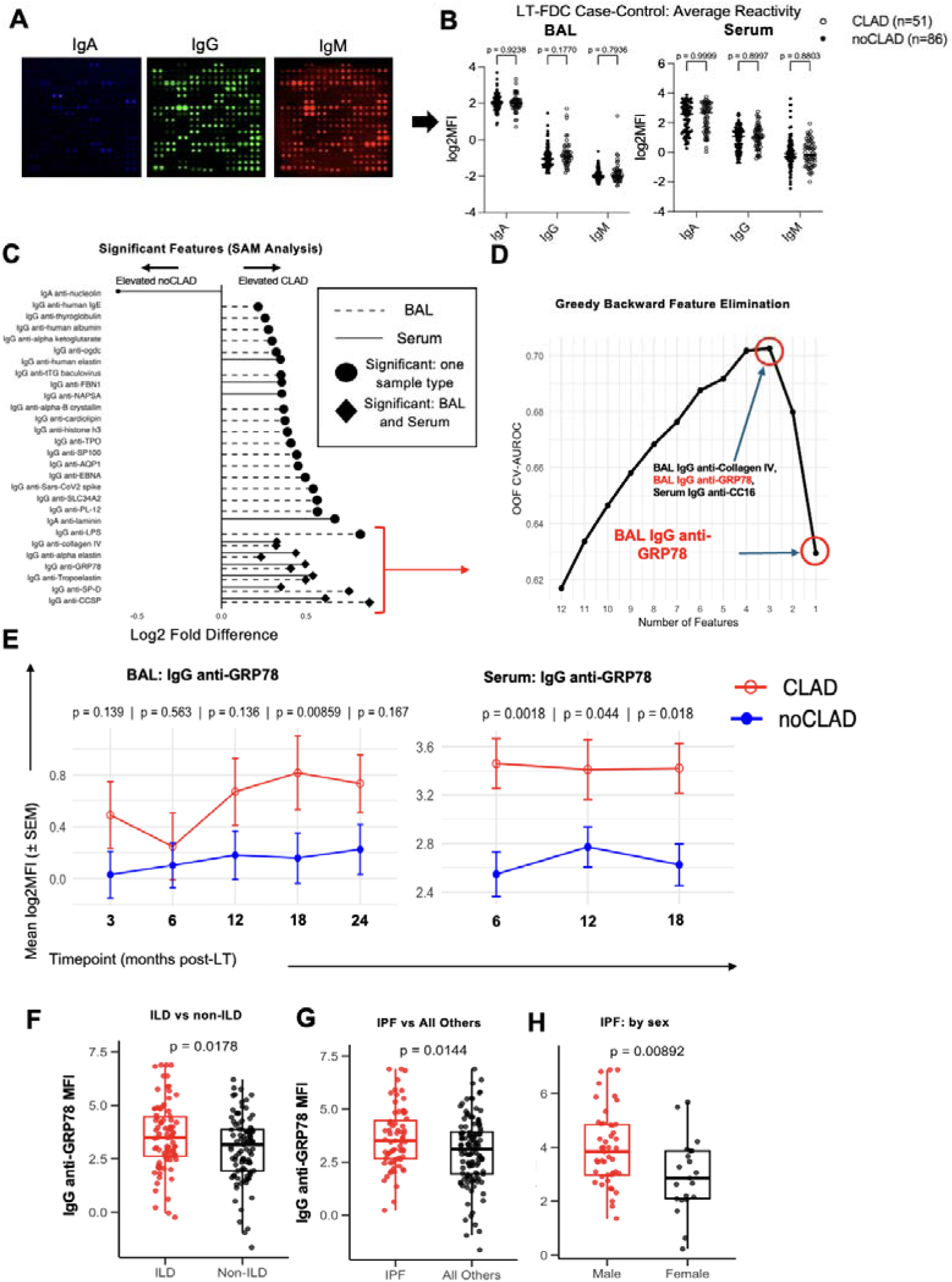
Antibody screening revealed IgG anti-GRP78 autoantibodies as associated with future CLAD and fibrotic lung diseases. **(A)** Representative microarray images demonstrating multiplexed detection of IgA, IgG, and IgM antibodies against a custom panel of antigens printed in duplicate. **(B)** Average serum and BAL antibody log_2_ mean fluorescent intensities (MFI) measured 5-7 months post-transplant in the LT-FDC cohort comparing patients who developed CLAD (open circles) within five years and those who remained CLAD-free (closed circles; cohort shown in Supplemental Figure 1A). **(C)** SAM analysis identifying antibodies significantly elevated in CLAD across serum and BAL samples (*q* < 0.05). **(D)** Combinatorial antibody scoring using cross-validated greedy backward stepwise selection restricted to antibodies elevated in both serum and BAL, showing feature selection and model performance across iterations. **(E)** Longitudinal BAL and serum IgG anti-GRP78 log_2_ MFI measured at multiple post-LT timepoints in patients case controlled for CLAD by five years. All samples were collected prior to CLAD diagnosis (Supplemental Figure 1B). **(F)** Serum IgG anti-GRP78 MFI across primary lung diseases. **(G)** Aggregate comparison of serum IgG anti-GRP78 MFI between ILDs and non-ILDs. **(H)** Serum IgG anti-GRP78 MFI in IPF compared with other lung diseases. **(I)** Serum IgG anti-GRP78 MFI in male and female patients with idiopathic pulmonary fibrosis.

To assess the temporal dynamics of anti-GRP78 IgG, in a longitudinal cohort case-controlled for CLAD by 5 years, BAL samples were collected at 3, 6-, 12-, 18-, and 24-months post-LT (Supplemental Figure 1B); for the same patients, serum samples were available at 6, 12, and 18 months post-LT. CLAD cases with CLAD onset less than 14 days following each time point were excluded to ensure relevance of measurements to future CLAD. Using our 64-pad microarray method (*20*), anti-GRP78 IgG levels in BAL progressively increased over the first two years post-LT in patients who later developed CLAD, notably rising around one-year post-LT, and serum reactivity remained consistently elevated across the measured timepoints (Figure 1E). Taken together, our findings reveal an early and persistent elevation in BAL IgG anti-GRP78 antibodies as a hallmark of patients developing CLAD within 5 years compared with patients who remained CLAD-free during this period.

### AT2 and transitional AT2 cells express HSPA5 in CLAD and IPF samples with RAS-specific enrichment

Detection of anti-GRP78 IgG antibodies in patients subsequently developing CLAD prompted us to investigate whether this antibody might also be detectable in ILD. Serum anti-GRP78 IgG levels were higher in interstitial lung diseases compard to non-ILDs (ILD; Figure 1F, *p* = 0.0178), especially in idiopathic pulmonary fibrosis (IPF; Figure 1G, *p* = 0.0144). IPF disproportionately affects men who suffer from significantly worse freedom from death and transplant than women (*33, 34*). Interestingly, males with IPF demonstrated significantly higher levels of anti-GRP78 IgG then women (Figure 1H, *p* = 0.00892).

Elevated anti-GRP78 IgG in patients with IPF and in LT recipients subsequently developing CLAD led us to investigate which pulmonary parenchymal cell populations exhibited elevated *HSPA5* expression (the gene encoding GRP78) using scRNA-seq. We generated data using fresh single-cell suspensions rapidly prepared from CLAD tissue immediately subjected to 3′ scRNA-seq. Excess donor lung (DL) that was obtained from a publicly available dataset generated at our institution was used as a comparator (*19*). Following data normalization, integration, and dimensionality reduction, epithelial cell clusters were identified and subsetted based on the canonical epithelial markers Epithelial Cell Adhesion Molecule (*EPCAM)* and E-cadherin (*CDH1)* (*35*), using Protein Tyrosine Phosphatase Receptor Type C (*PTPRC)* as an exclusion marker for hematopoietic lineage cells (Supplemental Figure 2). We identified 6 major clusters (Figure 2A) defined by canonical genes associated with distinct epithelial cell populations (Figure 2B). AT2 cells, characterized by surfactant protein C (*SFTPC)* and sodium-dependent phosphate transport protein 2B (*SLC34A2)* gene expression, organized into 3 clusters. Transitional AT2 cells can be further identified by the upregulation of canonical AT1-associated genes including Advanced Glycosylation End-Product Specific Receptor (*AGER)*, podoplanin (*PDPN)*, keratin 8 (*KRT8)* and claudin 4 (*CLDN4)* as described in IPF (*36–38*). A transitional identity was further supported by RNA velocity analysis (*39*) which indicated that transitional AT2 cells were positioned in the inferred trajectory downstream of AT2 cells and upstream of AT1 cells (Figure 2C). Of the remaining AT2 clusters, differential gene expression analysis suggested two AT2 programs, one surfactant-enriched (AT2 Active) and the other enriched for structural gene expression such as *TTN* and *MACF1* (AT2 Homeostatic; Supplemental Table 3). Notably, CLAD samples show a marked reduction in alveolar AT2 and AT1 cells in both proportion and number, suggesting a shift away from normal alveolar cellularity (Supplemental Figure 4 A-B).

**Figure 2.**
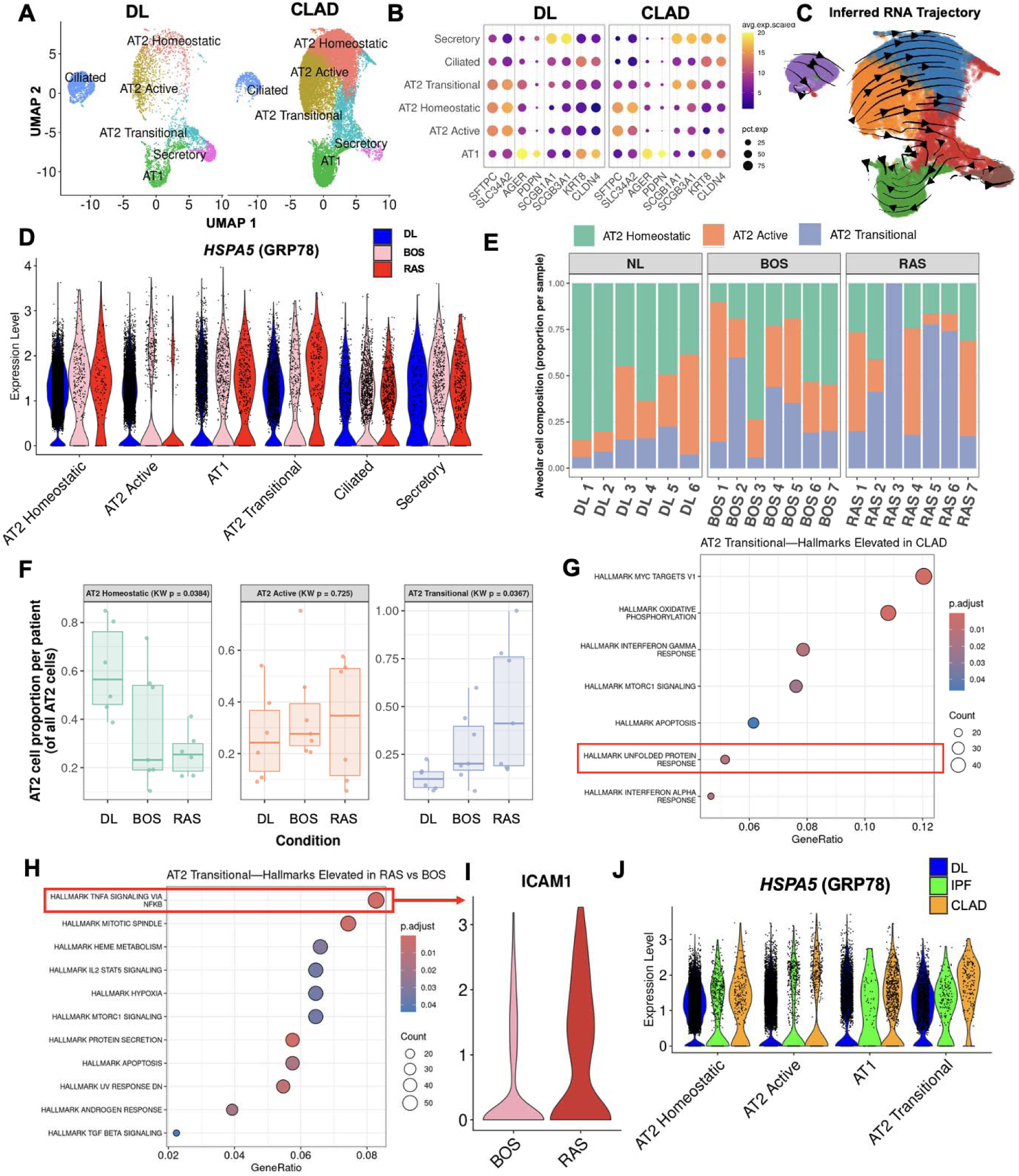
scRNA-seq identifies epithelial cell populations expressing *HSPA5* (GRP78) in CLAD and IPF. **(A)** UMAP embedding of epithelial cells from donor lungs (DL) and explanted CLAD tissue. **(B)**Dot plot showing expression of canonical genes associated with distinct epithelial cell lineages. **(C)**Inferred RNA velocity vectors projected onto the UMAP embedding. **(D)** *HSPA5* expression across epithelial cell clusters in DL, BOS, and RAS. **(E-F)** Distribution of AT2 cell proportions across DL, BOS, and RAS samples. **(G)** Pathway enrichment analysis of genes upregulated in transitional AT2 cells in CLAD compared to DL. **(H)** Comparison of pathway enrichment between BOS and RAS transitional AT2 cells. **(I)** Expression of *ICAM1* in transitional AT2 cells in BOS and RAS. **(J)** Integration of IPF scRNA-seq data with DL and CLAD samples subsetted for alveolar epithelial cells, with *HSPA5* expression shown across conditions.

*HSPA5* expression was increased in Active and Transitional AT2 cells in BOS and RAS compared with DL tissue and further increased in RAS relative to BOS in Transitional AT2 cells (Figure 2D). Similarly, there were more Transitional AT2 cells in CLAD compared to DL, particularly in RAS, mirroring IPF (*38, 40*) (Figure 2E-F). Differential gene expression analysis (Supplemental Figure 3A) revealed upregulation of the unfolded protein response (UPR) as a major pathway activated in Transitional AT2 cells in CLAD (Figure 2G). Other pathways significantly upregulated in CLAD Transitional AT2 cells may reflect biological processes that contribute to the UPR and subsequent *HSPA5* upregulation. For instance, IFN-γ – central to allograft rejection – can induce the UPR and GRP78 expression (*41, 42*). Comparing BOS and RAS (Supplemental Figure 3B) identified Hallmark Tumor Necrosis Factor (TNF)-α signaling via nuclear factor-κB (NF-κB) as the top upregulated pathway in RAS transitional AT2 cells (Figure 2H). Interestingly, TNF-α has been reported to upregulate GRP78 protein expression in primary synovial cells (*43*). Intercellular Adhesion Molecule 1 (*ICAM1)* was among the genes contributing to the identification of this pathway and may represent an important cell-surface marker for transitional AT2 cells in an aberrant fibrotic context (Figure 2I). Publicly available scRNA-seq data from 4 IPF lung samples (*44*) were integrated with our dataset and subsetted for alveolar epithelial cells. *HSPA5* expression was elevated in AT2 cells in both IPF and CLAD compared with DL lungs (Figure 2J). Together, these findings suggest that the elevated gene expression of *HSPA5* and accumulation of Transitional AT2 cells may represent a common and potentially central feature of fibrotic lung disease and processes involved in CLAD pathogenesis.

### Cell surface GRP78 expression is preferentially elevated on CLAD AT2 cells

We hypothesized that amplified *HSPA5* gene expression in CLAD-specific AT2 cells would correspond with increased AT2 cell-specific csGRP78 protein expression. Explanted peripheral lung tissue biopsies were obtained from five consecutive patients undergoing re-transplantation for CLAD, and five consecutive DL samples that were processed using the protocol described by Fujino et al. (*45*) (Figure 3A). This method was chosen for its superior epithelial cell proportional yield (*46*). Samples were subjected to analytical flow cytometry to assess csGRP78 expression on major populations (Figure 3B): all epithelial cells (CD45^-^CD31^-^EpCAM^+^), AT2 cells (CD45^-^CD31^-^EpCAM^-^HTII-280^+^), and club cells (CD45^-^CD31^-^HTII-280^-^ CCSP^high^). While endothelial cells, all epithelial cells, and club cells showed no significant differences in csGRP78 expression between CLAD and DL tissue, AT2 cells from CLAD lung tissue exhibited a striking increase in the proportion csGRP78^+^ AT2 cells (Figure 3C). Given the scarcity of re-transplantation cases, especially of a RAS or mixed CLAD phenotype, archival single-cell suspensions from patients with BOS (n = 5), RAS (n = 7), IPF (n = 11), and COPD (n = 6) were examined; these had been processed using the ColShort protocol described by Duong et al. (Figure 3A) known to yield proportionately fewer epithelial cells (Figure 3D) (*46*). AT2 csGRP78^+^ cells were elevated in BOS, RAS/mixed, and IPF, but not COPD, with respect to DL samples (Figure 3E). AT2 cells double-positive for csGRP78 and ICAM1 showed increased abundance in BOS, with greater enrichment in RAS and IPF (Figure 3E), consistent with transcriptional data indicating higher *ICAM1* expression in RAS AT2 cells (Figure 2H-I). Taken together, our data show that AT2 cells in CLAD and IPF exhibited elevated surface csGRP78 expression and were frequently double-positive for csGRP78 and ICAM1, particularly in RAS and IPF. These findings support the notion that AT2 ER stress may be a central feature of fibrotic lung disease, which may result in the formation of anti-GRP78 IgG.

**Figure 3.**
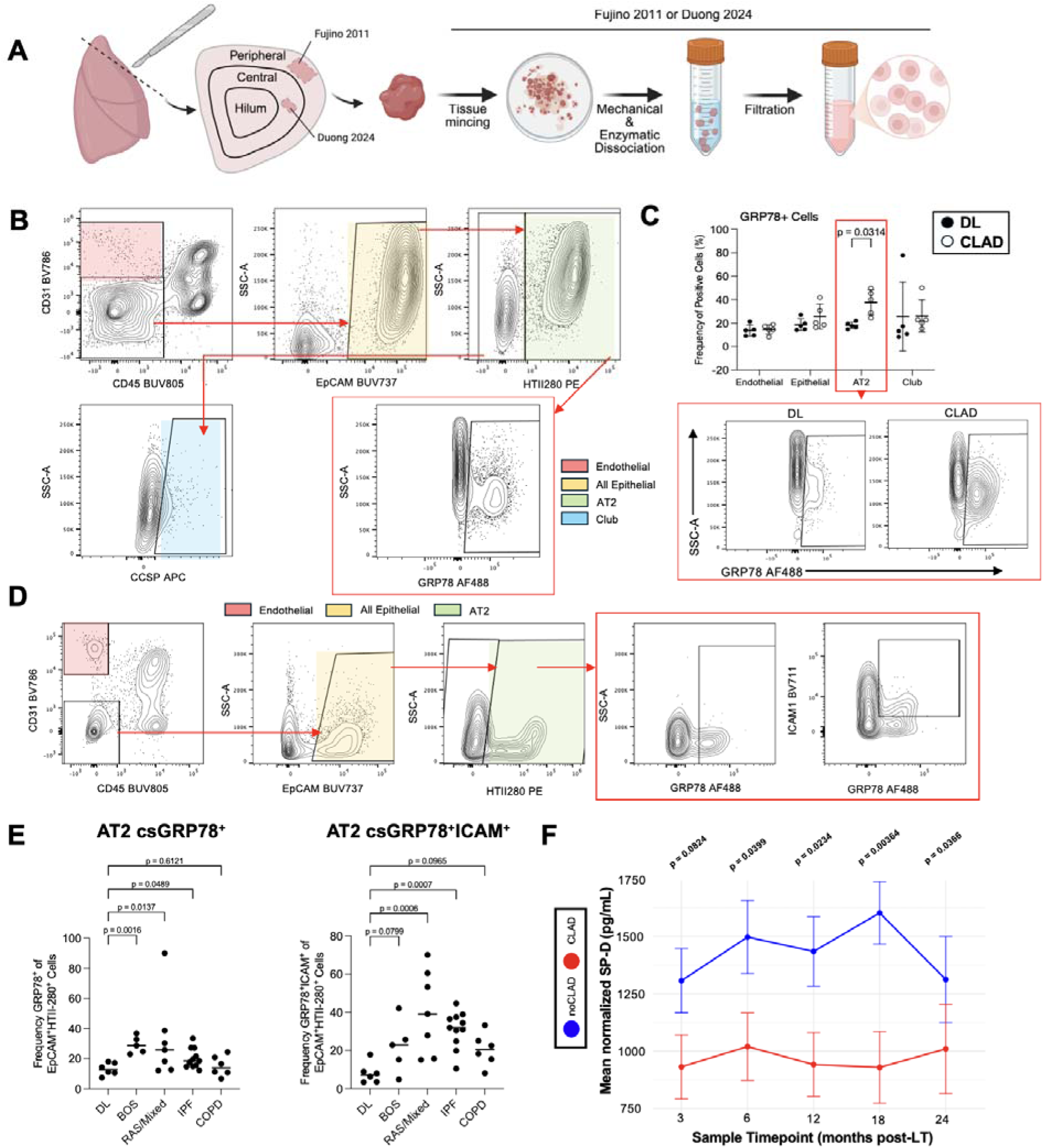
Flow cytometric assessment of csGRP78 on lung epithelial cell populations. **(A)**Schematic of lung tissue processing workflows used to generate single-cell suspensions for flow cytometry. **(B)** Gating strategy using the Fujino method to identify major lung cell populations including endothelial, total epithelial, AT2, and club cells. **(C)** Proportion of csGRP78^+^ cells in each population in DL and CLAD lung tissue. **(D)** Gating strategy for cells assessed using archival samples obtained with the ColShort lung tissue dissociation workflow. **(E)** Proportion of csGRP78^+^ AT2 cells and csGRP78^+^ICAM1^+^ AT2 cells across DL, BOS, RAS/mixed, IPF, and COPD. **(F)** BAL SP-D levels normalized to total protein concentration measured across longitudinal post-transplant timepoints in patients who developed CLAD and those who remained CLAD-free (cohort details shown in Supplemental Figure 1B). Panel **(A)** was made with Biorender.com.

In murine AT2 cells, ER stress has been shown to induce GRP78 and concurrently suppress surfactant protein transcription and content (*47*). If this pathway contributes to CLAD, ER stress-associated AT2 cell dysfunction could manifest as reduced alveolar surfactant protein levels detectable before clinical disease onset. In keeping with this possibility, BAL SP-D was consistently lower at every measured time point in those who developed CLAD compared with those who remained CLAD-free at 5 years (Figure 4A). Since this deficit was already present in the earliest samples we analyzed, it may indicate that AT2 cell loss or dysfunction begins well before CLAD is clinically apparent, perhaps as a result of injury in the perioperative period. We therefore hypothesized that perioperative storage and reperfusion serve as an initial priming event that triggers ER stress in AT2 cells, and that a failure to resolve this process may contribute to the pathogenesis of CLAD.

**Figure 4.**
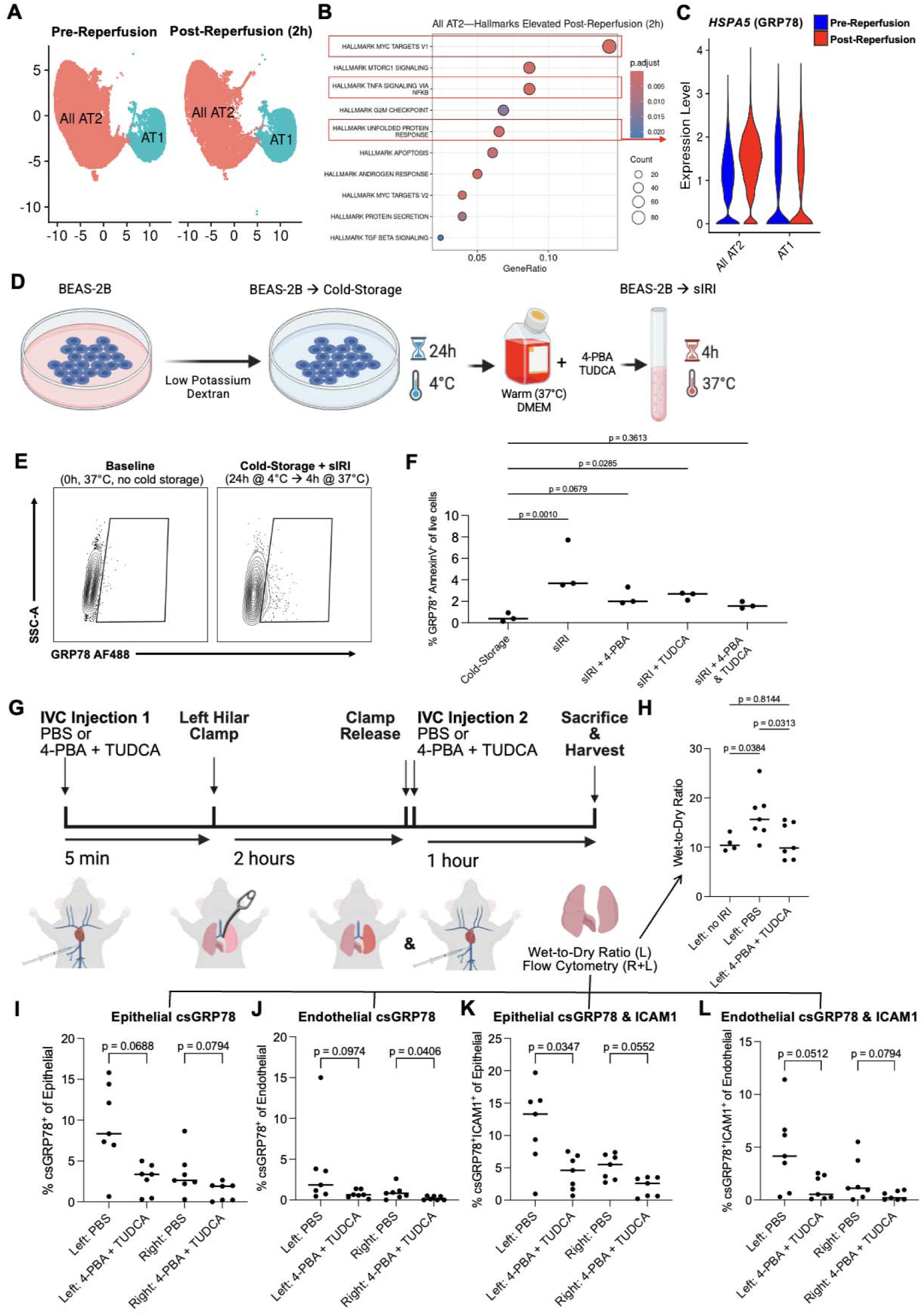
IRI-induced *HSPA5* and csGRP78 expression in lung epithelial and endothelial cells is attenuated by 4-PBA and TUDCA treatment. (A) UMAP embedding of alveolar epithelial cells from paired pre-and post-reperfusion lung biopsies generated using publicly available scRNA-seq data (*19*) and subsetted using canonical AT1 and AT2 markers. **(B)** Hallmark pathway enrichment analysis of genes differentially expressed between pre-and post-reperfusion samples. (C) *HSPA5* expression in AT2 and AT1 cells pre-and 2 hours post-reperfusion. **(D)** Schematic of the sIRI model using BEAS-2B cells. **(E)** csGRP78 expression following sIRI compared with cells maintained at 37 °C throughout. **(F)** csGRP78 expression following sIRI after cold-storage with addition of 4-PBA, TUDCA, or both to the reperfusion medium. **(G)** Schematic of the C57BL/10 murine left hilar clamp model of IRI. **(H)** Wet-to-dry ratio of left lungs from 4-PBA-TUDCA-treated, PBS-treated, and no-IRI animals. (**I-J**) csGRP78 expression in clamped left and contralateral right lung digests among **(I)** epithelial and **(J)** endothelial cells. **(K-L)** Proportion of csGRP78^+^ICAM-1^+^ cells in clamped left and contralateral right lung digests among **(K)** epithelial and **(L)** endothelial cells. Schematics in **(D)** and **(G)** were made with Biorender.com.

### IRI induces HSPA5 and cell surface GRP78 expression in human lung epithelial cells and is modulated by 4-PBA and TUDCA

To determine whether ER stress is engaged in AT2 cells during the transplant procedure itself, we analyzed publicly available scRNA-seq data from paired pre- and post-reperfusion lung biopsies from the Toronto Lung Transplant Program (*19*). Alveolar epithelial cells were subsetted using canonical AT2 and AT1 markers including *SFTPC*, *NAPSA*, *AGER*, and *PDPN* (Figure 4A). Hallmark analysis of differentially expressed genes revealed numerous pathways upregulated post-reperfusion, some of which were shared with CLAD (Figure 2G-H): MYC targets, TNFα signalling via NF-κB, and the UPR (Figure 4B). Within the UPR signature, *HSPA5* was upregulated at 2 hours post-reperfusion relative to cold ischemia in AT2 cells, with no corresponding increase seen in AT1 cells (Figure 4C).

To model cold storage and simulated ischemia-reperfusion injury (sIRI) in human lung epithelial cells, sub-confluent BEAS-2B cells were transferred to low-potassium dextran preservation solution at 4 °C for 24 h followed by return to DMEM with 10% FBS at 37 °C for 4 h (*48*) (Figure 4D). By analytical flow cytometry, csGRP78 was upregulated following sIRI relative to cells maintained at 37 °C throughout (no cold storage; Figure 4E). To determine whether this induction was modifiable, chemical chaperones were added during the reperfusion phase. Relative to media alone, 4-PBA and TUDCA each reduced csGRP78 expression when tested individually, and in combination (Figure 4F).

To assess whether 4-PBA and TUDCA might ameliorate IRI *in vivo*, we employed a mouse left hilar clamp model of IRI. The drugs were given in PBS by intravenous injection 5 minutes before hilar clamping and immediately after clamp release at 2 hours, prior to 1 hour of reperfusion (Figure 4G). Left lungs from 4-PBA-TUDCA-treated mice showed a reduction in wet-to-dry ratio compared with the PBS control group (Figure 4H), indicating a reduction in lung edema. Further, analytical flow cytometry demonstrated attenuated epithelial csGRP78 expression in the treated group; interestingly, a slight attenuation in epithelial csGRP78 expression in the contralateral right lung was also observed (Figure 4I). The same effect was observed in the endothelium (Figure 4J). Dual expression of csGRP78 and ICAM-1 also followed the same pattern in epithelium (Figure 4K) and endothelium (Figure 4K).

Together, these findings indicated that 4-PBA-TUDCA treatment mitigated pulmonary edema and attenuated markers of epithelial ER stress. The parallel endothelial changes implicated ER stress as a possible contributor to impaired endothelial integrity during IRI and suggested that mitigating endothelial stress may preserve the vascular barrier during the perioperative period. Notably, heightened expression of csGRP78 and ICAM-1 observed in the contralateral right lung raised the possibility that IRI in the left lung may have induced ER stress in tissue that was not rendered ischemic. If true, this may represent a mechanism by which injury sustained by one allograft could propagate to the contralateral lung. Systemically administered chemical chaperones would be positioned to act on both.

### Cell surface GRP78 expression is associated with fibrotic allograft pathology and is attenuated by 4-PBA-TUDCA treatment in an OLT model

IRI results in primary graft dysfunction (*49, 50*), and both are independently associated with CLAD (*51, 52*). How an injury sustained at reperfusion relates to dysfunction emerging months to years later has remained difficult to explain mechanistically. Our data point to one possible connection, in that the same cell surface phenotype appears in both settings. That this phenotype was pharmacologically reversible in hilar clamp IRI model raised the possibility that intervention against ER stress in AT2 cells beginning in the perioperative period may reduce the risk of CLAD following IRI. We tested this hypothesis using a minor alloantigen-mismatched C57BL/10-to-C57BL/6 (B10 B6) murine OLT model with IRI-augmented fibrotic pathology (*51*).

Recipients were given intraperitoneal 4-PBA-TUDCA (EXP) or PBS four times weekly from immediately after implantation until necropsy at 28-31 days post-OLT (Figure 5A). A composite left-lung pathology score (0–100) was calculated as the mean of twelve previously described histological parameters (*53*), each first normalized to its scale maximum so that every parameter contributed equally regardless of its native range. Composite scores were lower in EXP mice than PBS-treated mice (Figure 5C). Grafts were then stratified by median composite score. Epithelial csGRP78^+^ and csGRP78^+^ICAM-1^+^ cells were more frequent in grafts above the median, and epithelial csGRP78 correlated with composite score (Figure 5D). The same relationship was observed in endothelial cells (Figure 5E). EXP mice showed lower csGRP78 and csGRP78^+^ICAM-1^+^ frequencies in epithelial (Figure 5F) and endothelial cells (Figure 5G). Among individual histologic parameters, interstitial inflammation and parenchymal fibrosis were both lower in treated mice, as were ISHLT A-grade and vascular fibrosis, though none of these differences reached statistical significance (Figure 5H). Epithelial csGRP78 expression correlated significantly with each of these parameters (Figure 5I). Taken together, these data suggest that treatment with the chemical chaperones 4-PBA and TUDCA can mitigate epithelial and endothelial csGRP78 expression and CLAD-like pathology in a non-immunosuppressed mouse OLT model after IRI.

**Figure 5.**
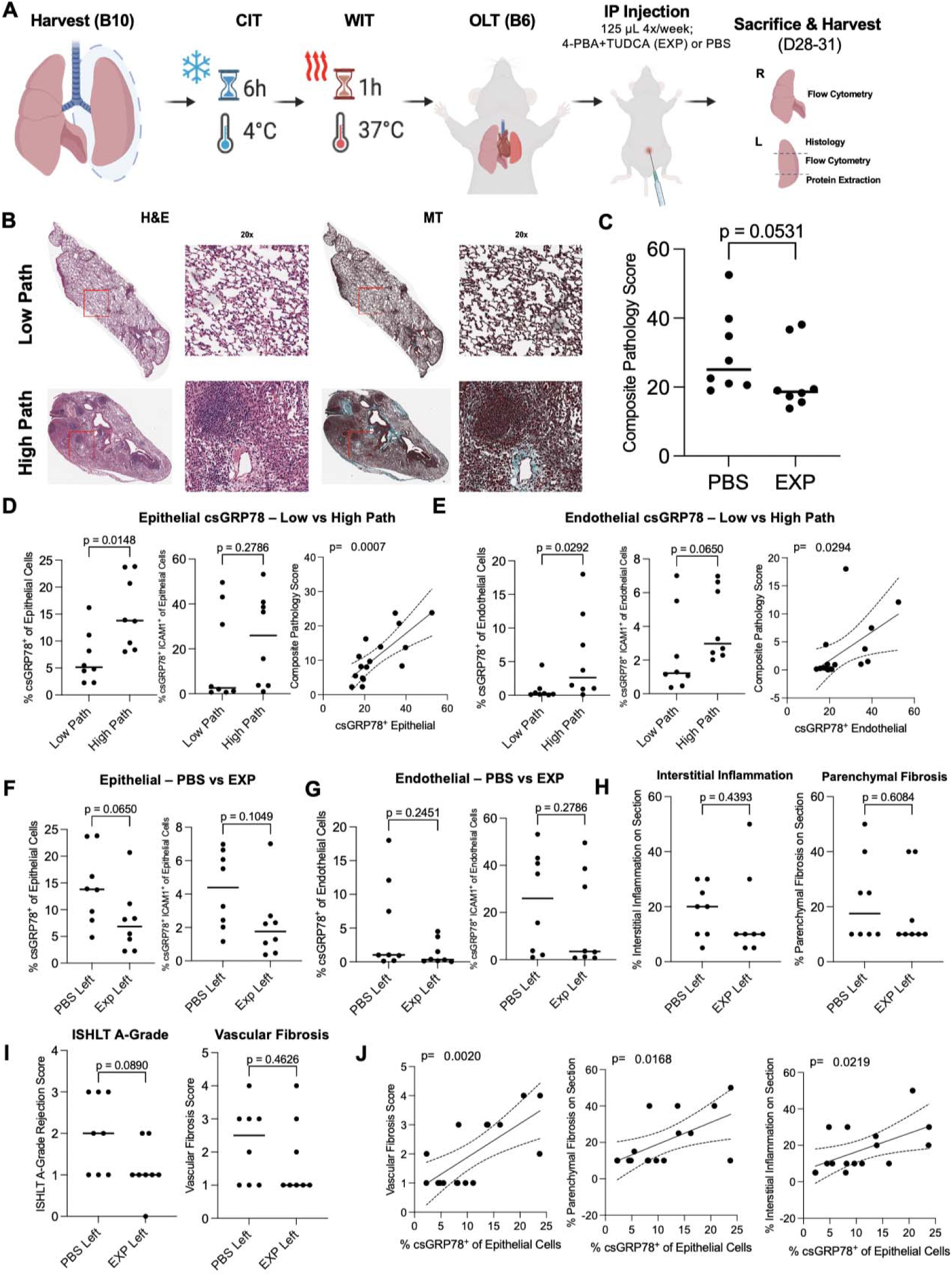
4-PBA and TUDCA treatment attenuates chronic allograft pathology and csGRP78 expression in a murine minor-mismatched OLT model. **(A)** Schematic of the B10 B6 OLT model and 4-PBA-TUDCA dosing schedule. **(B)** Representative hematoxylin and eosin [H&E] and Masson’s trichrome [MT] staining of left lung allografts from PBS-and 4-PBA-TUDCA-treated (EXP) mice. **(C)** Composite pathology score, calculated as the mean of twelve normalized histologic parameters, comparing EXP and PBS mice. **(D-E)** Proportion of csGRP78^+^ and csGRP78^+^ICAM-1^+^ epithelial **(D)** and endothelial **(E)** cells in grafts stratified by median composite score (left), with corresponding correlations against composite score (right). **(F-G)** Proportion of csGRP78^+^ and csGRP78^+^ICAM-1^+^ epithelial **(F)** and endothelial **(G)** cells comparing EXP and PBS mice. **(H-I)** Individual histologic parameters, including interstitial inflammation and parenchymal fibrosis **(H)**, and ISHLT A-grade and vascular fibrosis **(I)**, comparing EXP and PBS mice. **(I)** Relationship between csGRP78^+^ epithelial cells and individual histologic parameters. **(A)** was made with Biorender.com.

## DISCUSSION

Autoantibodies represent an increasingly appreciated facet of humoral immunity in lung transplantation. Those directed against KAT and ColV are previously-described correlates of future CLAD (*11–13*), and our recent identification of additional autoantibodies against lung-restricted targets further supports this paradigm (*32*). These findings indicate that self-tolerance breaks down after transplantation, likely sustained by the persistent inflammatory milieu driving CLAD. The numerous insults of the perioperative period may desequester self-antigens ordinarily subject to immunological ignorance. Presentation of such antigens in an inflammatory context might then undermine lung-specific self-tolerance. Beyond any direct pathogenic effect, these autoantibodies may serve as biomarkers of future rejection while reflecting the cellular processes from which their targets derive – a dimension that transplant autoantibody studies have seldom examined, and one that this work set out to address.

We uncovered autoantibodies in both serum and BAL shortly after transplantation that were associated with CLAD by 5 years. Greedy backward selection identified anti-GRP78 IgG as the most discriminative of these. BAL anti-GRP78 IgG rose progressively over the first two years after transplantation preceding CLAD while serum levels were elevated considerably earlier. This offset suggested compartment-specific kinetics, where a response arising in the periphery may later appear in the graft as local inflammatory conditions evolve. Anti-GRP78 IgG was also elevated in patients with ILD, and to a greater degree in IPF, suggesting that the process it reports is not unique to transplantation but shared among fibrotic lung diseases.

Our scRNA-seq analysis revealed increased *HSPA5* transcript within CLAD AT2 cells that was most prominent in Active and Transitional AT2 populations. Both populations were overrepresented in CLAD – the latter particularly in RAS. Interestingly, IFNγ, a dominant cytokine reflecting type 1 T cell-mediated allo-immunity in CLAD (*54, 55*), was upregulated in CLAD and has been shown to specifically induce the UPR and GRP78 expression in conjunctival secretory epithelial cells and in murine pulmonary epithelial cells (*41, 42*). Transitional AT2 cells showed signatures of UPR activation alongside TNFα/NF-κB signalling, a pathway implicated in driving GRP78 expression. TNFα neutralization has been shown to reduce ER stress both *in vitro* and *in vivo*, and conditioned media from TNFα-producing macrophages induced GRP78 in intestinal organoids (*56*). TNFα-induced ER stress in corneal epithelium was shown to be mitigated by 4-PBA and TUDCA (*57*). Together, these findings support a model in which inflammatory cues, particularly IFN-γ and TNF-α, may drive ER stress and GRP78 upregulation in CLAD epithelium. *ICAM1* was among the upregulated genes in this Hallmark pathway, possibly serving as a candidate surface marker of Transitional AT2 cells in a fibrotic context, and one that may perpetuate local inflammation through adhesion to LFA-1 on activated lymphocytes (*58*). Integration with existing IPF datasets revealed comparable *HSPA5* elevation within AT2 populations, suggesting that GRP78 upregulation and the accumulation of aberrant transitional epithelial states are shared features of fibrotic lung disease. It is important to note that the UPR, and the ensuing csGRP78 expression, may reflect an attempt at surviving inflammatory imjury and cell death – which may explain the reduced epithelial cell counts and proportions we observed.

Cell-surface translocation of GRP78 under ER stress, and its association with anti-GRP78 IgG, is well established across conditions including myelin oligodendrocyte glycoprotein antibody-associated disease, type 1 diabetes, cancer, atherosclerosis, and neuromyelitis optica (*21–30, 58*). Our findings indicate that AT2 epithelial cells from CLAD lungs are no exception: csGRP78^+^ AT2 cells were markedly increased in CLAD, with no corresponding statistically significant differences in other parenchymal populations. csGRP78^+^ AT2 cells co-expressing csGRP78 and ICAM-1 were particularly enriched in RAS and mixed CLAD, matching the transcriptional signature of transitional AT2 cells, and were also increased in IPF.

If csGRP78 marks AT2 cells under ER stress, that stress would be expected to carry a functional consequence. We therefore assessed BAL SP-D as a surrogate of AT2 secretory output, finding consistently lower levels across two years of routine follow-up in patients who subsequently developed CLAD. Reduced BAL SP-D following LT may therefore reflect a shift of the AT2 compartment out of its homeostatic secretory state, detectable well before fibrotic remodelling is established, and possibly as early as the occurrence of IRI at implantation. It may also reflect a specific loss of AT2 cells over time, which we observed at the time of explant.

Ischemia, an unavoidable consequence of organ procurement, depletes ATP and collapses the ion gradients maintained by energy-dependent transporters (*59*). The ER is particularly vulnerable: SERCA-mediated Ca^2+^ sequestration fails, and because the resident folding machinery is both Ca^2+^- and ATP-dependent, the luminal environment required for protein folding is lost (*60, 61*). Misfolded protein accumulates and sequesters GRP78 from PERK, IRE1α and ATF6, activating the UPR (*62*). Sustained UPR signalling has effects beyond the ER. PERK-driven ATF4 induces CHOP and, in turn, the ER oxidoreductase ERO1α that activates the inositol trisphosphate receptor (IP3R) to trigger Ca^2+^ efflux (*63*). At mitochondria-associated ER membranes, this Ca^2+^ can be delivered directly into the mitochondrial matrix (*64*). Reperfusion then supplies the oxygen for a ROS burst: oxidation of ischemia-accumulated succinate driving reverse electron transport at Complex I (*65*), xanthine oxidase acting on accumulated purine catabolites, and NADPH oxidase in recruited neutrophils (*66, 67*). Supraphysiological Ca^2+^ levels, ROS accumulation, and ATP depletion triggers the mitochondrial permeability transition pore, which is a mechanism of cell death (*68, 69*). ROS can oxidise nascent proteins and further activate IP3R (*70*). Ischemia may therefore initiate ER stress that can be amplified with reperfusion – Ca^2+^ carrying injury from ER to mitochondrion and ROS carrying it back in a potentially self-reinforcing circuit capable of sustaining dysfunction beyond the initial insult.

Several of our observations place the initiating event at reperfusion itself. In perioperative human samples, AT2 cells upregulated *HSPA5* within two hours of reperfusion relative to cold ischemia. *In vitro*, lung epithelial cells subjected to sIRI displayed csGRP78 at the surface, establishing that the ischemia-reperfusion sequence is sufficient to produce the phenotype, and chemical chaperone treatment during the reperfusion phase attenuated this induction. The same held *in vivo*: in a murine hilar clamp model, dual 4-PBA-TUDCA reduced wet-to-dry ratio while attenuating csGRP78 on both epithelium and endothelium. Together with reduced BAL SP-D early post-transplant, these data support the possibility of AT2 dysfunction that may be initiated by IRI and pharmacologically tractable.

Similarly, our data suggest that targeting the UPR in AT2 cells can limit fibrosis resulting from chronic rejection. In a minor-mismatched OLT model, csGRP78 tracked with graft pathology, sustained 4-PBA-TUDCA administration lowered csGRP78 and csGRP78 □ICAM-□1 frequencies in epithelium and endothelium alike, confirming that the target remains engageable across the chronic phase. These observations are significant, because conventional immunosuppressive therapies used in lung transplantation typically fail to limit fibrosis progression. Our findings suggest a complementary approach in which maladaptive cellular responses to injury are directly targeted to reduce fibrosis progression.

Our study has several limitations. We established associations between anti-GRP78 IgG, csGRP78 expression, and future CLAD, but not causality. Additional human cohorts will be required for external validation. Longitudinal sampling did not capture the perioperative period, with the earliest samples collected at 3 months post-transplant. SP-D served as an indirect surrogate for AT2 function without serial tissue validation. Severity of IRI, infectious exposure, and cross-reactivity or molecular mimicry could not be accounted for. Moreover, mechanisms governing surface GRP78 translocation were not directly tested. Trajectory inferences from scRNA-seq remain speculative in the absence of definitive lineage tracing. More broadly, simplified *in vitro* and murine systems cannot fully recapitulate the multicellular complexity of human CLAD, thus precluding firm conclusions about how these mechanisms would operate in the transplanted human lung.

In conclusion, anti-GRP78 IgG arose before CLAD onset and was similarly elevated in IPF. This pointed to a process shared across fibrotic lung disease. Its antigen was displayed at the surface of AT2 cells within hours of reperfusion post-LT to possibly position the origin of a chronic autoimmune response in the first moments of the transplant itself. That this phenotype proved pharmacologically reversible at reperfusion and remained targetable thereafter identifies AT2 cell ER stress as a tractable pathway from the perioperative window onward.

## MATERIALS AND METHODS

### Study design

To identify variable autoantibodies between patients who developed CLAD and those who remained free of CLAD by 5 years post-LT, we established a retrospective cohort of adults who underwent LT between September 2010 and December 2014 (Supplemental Figure 1A). Patients were classified in a case-control design as previously described (*32*). Briefly, single lung, multiorgan, and re-transplant patients were excluded to yield patients undergoing first bilateral LT. Patients who died before developing CLAD, lacked sufficient PFTs prior to sample collection, or underwent re-transplant without developing CLAD within the first 5 years post-LT were excluded. Patients with sera available within 6 months pre-LT and 5-to-7 months post-LT were included, and BAL at the post-LT timepoint was collected when available. This cohort was used to facilitate a larger screen of relevant features.

A longitudinal cohort was developed using samples collected during routine follow up at 3-, 6-, 12-, 18-, and 24-months post-LT, requiring BAL availability at these timepoints. Serum was collected at 6, 12, and 18 months, with samples matched to the same patients and timepoints as BAL (Supplemental Figure 1B). Similar patient selection criteria were applied, such that samples at these timepoints were collected prior to CLAD diagnosis, and patients were case controlled for future CLAD by 5 years. All serum samples were obtained from the University Health Network’s Histocompatibility Laboratory, and BAL samples were obtained from the Toronto Lung Transplant Program’s Biobank. Demographic variables for both cohorts are presented in Supplemental Table 2. The study was approved by the University Health Network’s Research Ethics Board (15-9531).

### Antigen microarray

Antibody detection was performed using our custom antigen microarray protocol (*20*). Briefly, antigens prepared to a final concentration of 200 mg/mL were spotted onto 2-pad or 64-pad nitrocellulose-coated glass slides on the ONCYTE® SuperNOVA platform (Grace Bio-Labs, Bend, OR, USA). After blocking with microarray blocking buffer (2% BSA, 7.5% fetal bovine serum, and 0.1% Tween-20 in PBS), BAL or serum samples were diluted 1:1 or 1:100, respectively, in microarray blocking buffer and applied to microarray membranes using incubation chambers (2-pad: GVS FAST™ incubation chambers, Maine Manufacturing, ME, USA; 64-pad: ProPlate® 64 Well Slide Module, Grace Bio-Labs, Bend, OR, USA). Secondary detection antibodies were prepared in microarray blocking buffer at final dilutions of 1:1000 goat anti-human IgM-Cy5 (polyclonal, Jackson ImmunoResearch; 115-175-075), 1:1000 goat anti-human IgA-fluorescein (FITC; polyclonal, ThermoFisher; A18782), and 1:2000 goat anti-human IgG-Cy3 (polyclonal, Jackson ImmunoResearch; 109-165-003). Median fluorescence intensities (MFI) of antibody reactivity were quantified using GenePix Pro (software version 6.1.0.2, Molecular Devices, Sunnyvale, CA).

### Lung tissue collection and processing

Explanted CLAD lung tissue was collected at the time of re-transplantation. Samples were obtained from the right upper lobe. Single-cell suspensions of central tissue were rapidly generated using the ColShort protocol described by Duong et al. (*46*), using approximately 1 x 2 x 2 cm tissue sections, and were immediately processed for 3′ scRNA-seq on the 10x platform. Additional aliquots were resuspended in freezing media (90% FBS, 10% DMSO) and cryopreserved. Samples generated using the Fujino et al. protocol (*45*), which provides superior proportional epithelial cell yield (*46*), were prepared from approximately 3g peripheral right upper lobe tissue sections, with aliquots similarly cryopreserved for future use.

### Single-cell RNA sequencing data analysis

CLAD samples were subject to 3′ scRNA-seq and subsequently analyzed in R using Seurat. Integration with publicly available DL (*19*) and IPF (*44*) was performed and subsetted for epithelial cells. Differential expression, Hallmark pathway enrichment, and RNA velocity (*39*) analytical methodology are described Supplementary Materials and Methods.

### Flow cytometry

Cryopreserved single-cell suspensions of human lung tissue were rapidly thawed in a 37°C water bath then washed and pelleted before resuspension in Phosphate-Buffered Saline (PBS). Samples were washed with PBS and stained with BD Horizon™ Fixable Viability Stain 575V (1:500; Becton, Dickinson and Company [BD]; 565694) in PBS for 12 minutes at room temperature, then washed with flow cytometry staining buffer (FCS; PBS, 2% FBS, 2 mM EDTA) and pelleted prior to surface staining. All subsequent washes were performed in FCS with centrifugation at 400 x g for 5 minutes and supernatant removal.

Epithelial cells were stained using a three-step approach optimized for AT2 cell detection based on Eiliazadeh et al. (*71*). First, samples were incubated with 50 µL of mouse IgM anti-human HTII-280 primary antibody (1:10; Terrace Biotech: TB-27AHT2-280) for 15 minutes at 4°C, washed, and then incubated with 50 µL of PE-conjugated anti-mouse IgM secondary antibody (1:200; BioLegend: 406508) for 10 minutes at 4°C. Samples were then washed and stained with 50 µL of a cocktail of antibodies in FCS to delineate parenchymal cell populations as described in this study: CD45 (BUV805, 1:100; BD Horizon™: 612891), CD31 (BV786, 1:100; BD OptiBuild™: 744467), EpCAM/CD326 (BUV737, 1:100; BD OptiBuild™: 748382), GRP78 (Alexa Fluor™ 488, 1:100; Invitrogen™: 53976880), CD200 (BB700, 1:100; BD Horizon™: 566536), CD148 (BUV661, 1:100; BD OptiBuild™: 750009), and CD54/ICAM-1 (BV711, 1:200; BD Horizon™: 564078). Samples were washed and resuspended in 200 µL of 4% paraformaldehyde (PI28908, Thermo Fisher Scientific) in PBS for 20 minutes at 4 °C. Cells were washed and resuspended in 250 µL of FCS for flow cytometric analysis on the BD FACSymphony™ A5 SE Cell Analyzer. Flow cytometry data were analyzed using FlowJo software (v10.10; Tree Star, San Carlos, CA, USA).

### Surfactant protein D ELISA and total protein quantification

BAL samples were collected from the longitudinal follow-up cohort (Supplemental Figure 1B), and SP-D levels were measured using the Human SP-D DuoSet ELISA kit at a dilution of 1:50 (R&D Systems; DY1920). BAL SP-D concentrations were normalized to total protein to account for variability in sample collection (*72*). Total protein concentration was measured using the Pierce bicinchoninic acid (BCA) Protein Assay Kit (Thermo Fisher Scientific; 23227).

### In vitro model of cold storage and simulated ischemia-reperfusion injury

BEAS-2B cells were cultured in low-glucose Dulbecco’s Modified Eagle Medium (DMEM) supplemented with 10% fetal bovine serum (FBS; Gibco, 12484028) until confluence. To model cold storage and simulated ischemia-reperfusion injury (sIRI), a protocol adapted from Saren et al. was used (*48*). Culture medium was replaced with low-potassium dextran (LPD) solution and cells were stored at 4 °C for 24 hours. Cells were detached with trypsin-EDTA. For each replicate, a portion of cells was immediately stained for flow cytometry with Fixable Viability Stain 575V and anti-GRP78 (Alexa Fluor 488, 1:100; Invitrogen, 53976880), washed with 1x Annexin V binding buffer prepared in ddH_2_O (Thermo Fisher, V13246), and stained with 5 µL Annexin V (BUV395; BD Horizon, 564871) per sample in residual fluid for 20 minutes at 4 °C. Cells were then washed again in binding buffer and fixed in 4% PFA. The remaining cells from each replicate were subjected to sIRI by replacing LPD with DMEM supplemented with 10% FBS prewarmed to 37 °C. Where indicated, the reperfusion medium was further supplemented with 10 mM sodium 4-phenylbutyrate (4-PBA; MedChemExpress, HY-15654) and/or tauroursodeoxycholic acid sodium salt (TUDCA; MedChemExpress, HY-19696A). Cells were incubated at 37 °C with 5% atmospheric CO_2_ for 4 hours, stained as above, and analyzed on a BD FACSymphony A5 SE Cell Analyzer.

### Mouse hilar clamp model of ischemia-reperfusion injury

Animal procedures were approved by the institutional animal care committee and conducted in accordance with the Canadian Council on Animal Care guidelines (Animal Use Protocol 7310). Male C57BL/10 (B10) mice received 125 µL of 4-PBA (100 mg/kg) and TUDCA (250 mg/kg) in PBS (EXP) or an equal volume of PBS (control) by inferior vena cava injection 5 minutes before clamping. The left hilum was occluded with an aneurysm clip for 2 hours of warm ischemia, after which the clip was released and a second 125 µL dose administered immediately. Following 1 hour of reperfusion, mice were sacrificed and the left (clamped) and right (non-clamped) lungs harvested. Single-cell suspensions were prepared from both lungs seperately similar to ColShort methods from Duong et al (*46*). Briefly, lungs were digested in 4 mL of Hanks’ Balanced Salt Solution containing collagenase A (1 mg/mL) and DNase I (0.2 mg/mL) and mechanically dissociated using a gentleMACS tissue dissociator (Miltenyi Biotec, Bergisch Gladbach, Germany). The digest was filtered, subjected to red blood cell lysis, and resuspended for flow cytometric analysis. Washing and staining were performed as described for human samples, except that a single extracellular antibody cocktail was used: BUV737 rat anti-mouse CD45 (BD OptiBuild, 752414), BB700 rat anti-mouse CD31 (BD Horizon, 566490), APC/Cyanine7 anti-mouse CD326 (EpCAM; BioLegend, 118218), Alexa Fluor 488-conjugated polyclonal anti-GRP78 (Thermo Fisher, PA1-014A-A488), and Alexa Fluor 647 rat anti-mouse CD54 (ICAM-1; BioLegend, 116114).

### Minor alloantigen mismatched mouse model of chronic lung allograft rejection

Male C57BL/10 (B10, H-2^b^) donor lungs were transplanted into C57BL/6 (B6, H-2^b^) recipients after 6 hours of cold and 1 hour of warm ischemia, as previously described (*51*). Recipients received EXP or control solution intraperitoneally 4x/week until necropsy at 28-31 days. Left lung tissue was divided for flow cytometry, histology, and protein extraction. Additional details are provided in the Supplementary Materials and Methods.

### Mouse lung allograft pathology assessment

Pathology severity scoring was performed by a blinded observer (AP) using our published method (*53*). A composite pathology score (0–100) was determined for each graft as the mean of twelve previously described histological parameters, each first normalized to its scale maximum so that every parameter contributed equally regardless of its native range.

### Statistical analysis

Between-group analyses of microarray-determined antibody reactivities were compared using SAM (*32*) and two-way ANOVA with Šídák correction. Cross-validated backward elimination was used to identify important features using autoantibody composite scoring (*32*) and assessed by AUROC. Wilcoxon rank-sum, two-tailed t-tests, Kruskal-Wallis tests with Dunn’s post hoc testing, or linear regression was performed for other comparisions. R and GraphPad Prism was used for analyses. Software, versions, and further details are given in the Supplementary Materials and Methods.

## List of Supplementary Materials Supplemental Materials and Methods

**Fig. S1.** CONSORT diagrams depicting patient selection and cohort structure.

**Fig. S2.** Epithelial cell isolation from scRNA-seq data.

**Fig. S3.** Differential gene expression in transitional AT2 cells.

**Fig. S4.** Per-patient epithelial cell composition in DL vs CLAD.

**Supplemental Table S1.** Characteristics of the nested case-control subset and Time-to-CLAD cohort.

**Supplemental Table S2.** List of antigens used in this study.

**Supplemental Table S3.** Differential gene expression comparing AT2 Active and AT2 Homeostatic cell clusters in donor lungs.

## Acknowledgements

The authors wish to thank Marcelo Cuesta, Erika Kurtz, and Ella Arustei of the Toronto Lung Transplant Program biobank, and Dr. Jeffrey Kiernan of the UHN HLA Laboratory for assistance with procuring serum and BAL samples. We also wish to thank Rasheed Ghany and the Toronto Lung Transplant Program Database team for assistance with clinical data collection and curating. Schematic figures were made with Biorender.com.

## Funding

Canadian Institutes of Health Research Project Grant PJT173343 (SJ, TM)

International Society for Heart and Lung Transplantation/CareDx Non-invasive Biomarker Grant (SJ)

Caron Thorburn Institute/UHN Foundation (SJ, TM)

Truscott-Parkes Catalyst Fund in Idiopathic Pulmonary Fibrosis (SJ) DiPoce Research Scholar Award (TM)

Sanofi iAward (TM, SJ)

Canadian Society of Transplantation Research Training Award (DV)

Canadian Institutes of Health Research Canada Graduate Scholarship – Doctoral (DV)

Canadian Institutes of Health Research Canada Graduate Scholarship – Master’s (DV)

Natural Sciences and Engineering Research Council of Canada: Collaborative Research and Training Experience fellowship in Cell and Engineering Approaches to Preserve and Rejuvenate Organs (DV)

University of Toronto Open Fellowship, Institute of Medical Science, Temerty Faculty of Medicine (DV)

## Author Contributions

Conceptualization: DV, SJ

Methodology: DV, AC, AL, TA, AP, AD, KFB, ME, NC, TM, SJ

Investigation: DV, AC, AL, TA, AP, AD, KFB, TM, SJ

Visualization: DV, AL

Funding acquisition: SJ, TM, DV

Project administration: SJ, TM, SK, MC, ML

Supervision: SJ, TM, ML, DV

Writing – original draft: DV, SJ

Writing – review & editing: DV, AL, TA, AP, AD, KFB, AC, ML, SK, MC, TM, SJ

## Competing Interests

SJ and TM: Sanofi Inc. CLAD advisory board

SJ has received speaker’s fees from Therakos, Inc.

SJ, SK, DV and TM have applied for a US Patent (63/560,416) entitled, “AUTOANTIBODY PROFILING TO PREDICT CHRONIC LUNG ALLOGRAFT DYSFUNCTION” in relation to this work.

## Data and materials availability

All data are available on reasonable request. Materials can be made available subject to a materials transfer agreement.

## Notes

### Competing Interest Statement

Competing Interests:
SJ and TM: Sanofi Inc. CLAD advisory board
SJ has received speaker fees from Therakos Inc.
SJ SK DV and TM have applied for a US Patent (63/560416) in relation to this work.

## References

1. D. C. Chambers, W. S. Cherikh, M. O. Harhay, D. Hayes, E. Hsich, K. K. Khush, B. Meiser, L. Potena, J. W. Rossano, A. E. Toll, T. P. Singh, A. Sadavarte, A. Zuckermann, J. Stehlik, The International Thoracic Organ Transplant Registry of the International Society for Heart and Lung Transplantation: Thirty-sixth adult lung and heart–lung transplantation Report—2019; Focus theme: Donor and recipient size match. J. Heart Lung Transplant. 38, 1042–1055 (2019).

2. C. N. Graham, C. Watson, A. Barlev, M. Stevenson, V. R. Dharnidharka, Mean lifetime survival estimates following solid organ transplantation in the US and UK. J. Med. Econ. 25, 230–237 (2022).

3. M. Sato, T. K. Waddell, U. Wagnetz, D. M. Hwang, A. Haroon, D. Wagnetz, C. Chaparro, L. G. Singer, M. A. Hutcheon, S. Keshavjee, Restrictive allograft syndrome (RAS): a novel form of chronic lung allograft dysfunction. J. Heart Lung Transplant. Off. Publ. Int. Soc. Heart Transplant. 30, 735–742 (2011).

4. W. A. Wuyts, A. Cavazza, G. Rossi, F. Bonella, N. Sverzellati, P. Spagnolo, Differential diagnosis of usual interstitial pneumonia: when is it truly idiopathic? Eur. Respir. Rev. 23, 308– 319 (2014).

5. A. Adegunsoye, C. A. Newton, J. M. Oldham, B. Ley, C. T. Lee, A. L. Linderholm, J. H. Chung, N. Garcia, D. Zhang, R. Vij, R. Guzy, R. Jablonski, R. Bag, R. S. Voogt, S.-F. Ma, A. I. Sperling, G. Raghu, F. J. Martinez, M. E. Strek, P. J. Wolters, C. K. Garcia, B. L. Pierce, I. Noth, Telomere length associates with chronological age and mortality across racially diverse pulmonary fibrosis cohorts. Nat. Commun. 14, 1489 (2023).

6. S. Everaerts, E. J. Lammertyn, D. S. Martens, L. J. De Sadeleer, K. Maes, A. A. van Batenburg, R. Goldschmeding, C. H. M. van Moorsel, L. J. Dupont, W. A. Wuyts, R. Vos, G. Gayan-Ramirez, N. Kaminski, J. C. Hogg, W. Janssens, G. M. Verleden, T. S. Nawrot, S. E. Verleden, J. E. McDonough, B. M. Vanaudenaerde, The aging lung: tissue telomere shortening in health and disease. Respir. Res. 19, 95 (2018).

7. P.-J. Royer, G. Olivera-Botello, A. Koutsokera, J.-D. Aubert, E. Bernasconi, A. Tissot, C. Pison, L. Nicod, J.-P. Boissel, A. Magnan, on behalf of the S. Consortium, Chronic Lung Allograft Dysfunction: A Systematic Review of Mechanisms. Transplantation 100, 1803–1814 (2016).

8. M. Keller, S. Yang, L. Ponor, A. Bon, A. Cochrane, M. Philogene, E. Bush, P. Shah, J. Mathew, A. W. Brown, H. Kong, A. Charya, H. Luikart, S. D. Nathan, K. K. Khush, M. Jang, S. Agbor-Enoh, Preemptive treatment of *de novo* donor-specific antibodies in lung transplant patients reduces subsequent risk of chronic lung allograft dysfunction or death. Am. J. Transplant. 23, 559–564 (2023).

9. J. M. Tikkanen, L. G. Singer, S. J. Kim, Y. Li, M. Binnie, C. Chaparro, C.-W. Chow, T. Martinu, S. Azad, S. Keshavjee, K. Tinckam, De Novo DQ Donor-Specific Antibodies Are Associated with Chronic Lung Allograft Dysfunction after Lung Transplantation. Am. J. Respir. Crit. Care Med. 194, 596–606 (2016).

10. F. P. Legoux, J.-B. Lim, A. W. Cauley, S. Dikiy, J. Ertelt, T. J. Mariani, T. Sparwasser, S. S. Way, J. J. Moon, CD4+ T Cell Tolerance to Tissue-Restricted Self Antigens Is Mediated by Antigen-Specific Regulatory T Cells Rather Than Deletion. Immunity 43, 896–908 (2015).

11. U. Rao, M. Sharma, T. Mohanakumar, C. Ahn, A. Gao, V. Kaza, Prevalence of antibodies to lung self-antigens (Kα1 Tubulin and Collagen V) and Donor Specific Antibodies to HLA in Lung Transplant Recipients and Implications for Lung Transplant Outcomes: Single Center Experience. Transpl. Immunol. 54, 65–72 (2019).

12. Alloimmunity-induced autoimmunity as a potential mechanism in the pathogenesis of chronic rejection of human lung allografts -PubMed (available at https://pubmed.ncbi.nlm.nih.gov/21414808/).

13. A. McQuiston, A. Emtiazjoo, P. Angel, T. Machuca, J. Christie, C. Atkinson, Set Up for Failure: Pre-Existing Autoantibodies in Lung Transplant. Front. Immunol. 12, 711102 (2021).

14. T. Iwata, A. Philipovskiy, A. J. Fisher, R. G. Presson, M. Chiyo, J. Lee, E. Mickler, G. N. Smith, I. Petrache, D. B. Brand, W. J. Burlingham, B. Gopalakrishnan, D. S. Greenspan, J. D. Christie, D. S. Wilkes, Anti-Type V Collagen Humoral Immunity in Lung Transplant Primary Graft Dysfunction. J. Immunol. Baltim. Md 1950 181, 5738–5747 (2008).

15. S. Yoshida, A. Haque, T. Mizobuchi, T. Iwata, M. Chiyo, T. J. Webb, L. A. Baldridge, K. M. Heidler, O. W. Cummings, T. Fujisawa, J. S. Blum, D. D. Brand, D. S. Wilkes, Anti-Type V Collagen Lymphocytes that Express IL-17 and IL-23 Induce Rejection Pathology in Fresh and Well-Healed Lung Transplants. Am. J. Transplant. 6, 724–735 (2006).

16. A. Bharat, S. Chiu, Z. Zheng, H. Sun, A. Yeldandi, M. M. DeCamp, H. Perlman, G. R. S. Budinger, T. Mohanakumar, Lung-Restricted Antibodies Mediate Primary Graft Dysfunction and Prevent Allotolerance after Murine Lung Transplantation. Am. J. Respir. Cell Mol. Biol. 55, 532– 541 (2016).

17. K. Farahnak, Y. Z. Bai, Y. Yokoyama, D. B. Morkan, Z. Liu, J. M. Amrute, A. D. F. Falcon, Y. Terada, F. Liao, W. Li, H. M. Shepherd, R. R. Hachem, V. Puri, K. J. Lavine, A. E. Gelman, A. Bharat, D. Kreisel, R. G. Nava, B cells mediate lung ischemia/reperfusion injury by recruiting classical monocytes via synergistic B cell receptor/TLR4 signaling. J. Clin. Invest. 134 (2024), doi:10.1172/JCI170118.

18. B. A. Whitson, M. E. Prekker, C. S. Herrington, T. P. M. Whelan, D. M. Radosevich, M. I. Hertz, P. S. Dahlberg, Primary Graft Dysfunction and Long-term Pulmonary Function After Lung Transplantation. J. Heart Lung Transplant. 26, 1004–1011 (2007).

19. A. Wong, A. Duong, G. Wilson, J. Yeung, S. MacParland, H. Han, M. Cypel, S. Keshavjee, M. Liu, Ischemia-reperfusion responses in human lung transplants at the single-cell resolution. Am. J. Transplant. 24, 2199–2211 (2024).

20. D. Vosoughi, A. Chruscinski, O. L. Rojas, S. Keshavjee, T. Martinu, S. Juvet, A scalable three-colour antigen microarray for simultaneous IgM, IgG, and IgA profiling in serum and bronchoalveolar lavage samples in health and disease (2026).

21. A. Raiter, A. Vilkin, R. Gingold, Z. Levi, M. Halpern, Y. Niv, B. Hardy, The presence of anti-GRP78 antibodies in the serum of patients with colorectal carcinoma: a potential biomarker for early cancer detection. Int. J. Biol. Markers 29, e431–435 (2014).

22. M. Gonzalez-Gronow, U. Gopal, R. C. Austin, S. V. Pizzo, Glucose-regulated protein (GRP78) is an important cell surface receptor for viral invasion, cancers, and neurological disorders. IUBMB Life 73, 843–854 (2021).

23. F. Shimizu, R. Ogawa, Y. Mizukami, K. Watanabe, K. Hara, C. Kadono, T. Takahashi, T. Misu, Y. Takeshita, Y. Sano, M. Fujisawa, T. Maeda, I. Nakashima, K. Fujihara, T. Kanda, GRP78 Antibodies Are Associated With Blood-Brain Barrier Breakdown in Anti–Myelin Oligodendrocyte Glycoprotein Antibody–Associated Disorder. Neurol. Neuroimmunol. Neuroinflammation 9, e1038 (2021).

24. S. Vig, M. Buitinga, D. Rondas, I. Crèvecoeur, M. van Zandvoort, E. Waelkens, D. L. Eizirik, C. Gysemans, P. Baatsen, C. Mathieu, L. Overbergh, Cytokine-induced translocation of GRP78 to the plasma membrane triggers a pro-apoptotic feedback loop in pancreatic beta cells. Cell Death Dis. 10, 309 (2019).

25. M. Buitinga, A. Callebaut, F. Marques Câmara Sodré, I. Crèvecoeur, G. Blahnik-Fagan, M.-L. Yang, M. Bugliani, D. Arribas-Layton, M. Marré, D. P. Cook, E. Waelkens, R. Mallone, J. D. Piganelli, P. Marchetti, M. J. Mamula, R. Derua, E. A. James, C. Mathieu, L. Overbergh, Inflammation-Induced Citrullinated Glucose-Regulated Protein 78 Elicits Immune Responses in Human Type 1 Diabetes. Diabetes 67, 2337–2348 (2018).

26. G. Liu, J. Wu, Y. Wang, Y. Xu, C. Xu, G. Fang, X. Li, J. Chen, The Differential Expressions and Associations of Intracellular and Extracellular GRP78/Bip with Disease Activity and Progression in Rheumatoid Arthritis. Bioengineering 12, 58 (2025).

27. M. Wang, S. Wey, Y. Zhang, R. Ye, A. S. Lee, Role of the Unfolded Protein Response Regulator GRP78/BiP in Development, Cancer, and Neurological Disorders. Antioxid. Redox Signal. 11, 2307–2316 (2009).

28. A. O. Akinyemi, K. E. Simpson, S. F. Oyelere, M. Nur, C. M. Ngule, B. C. D. Owoyemi, V. A. Ayarick, F. F. Oyelami, O. Obaleye, D.-P. Esoe, X. Liu, Z. Li, Unveiling the dark side of glucose-regulated protein 78 (GRP78) in cancers and other human pathology: a systematic review. Mol. Med. 29, 112 (2023).

29. E. D. Crane, A. A. Al-Hashimi, J. Chen, E. G. Lynn, K. D. Won, Š. Lhoták, M. Naeim, K. Platko, P. Lebeau, J. H. Byun, B. Shayegan, J. C. Krepinsky, K. J. Rayner, S. Marchiò, R. Pasqualini, W. Arap, R. C. Austin, Anti-GRP78 autoantibodies induce endothelial cell activation and accelerate the development of atherosclerotic lesions. JCI Insight 3 (2018), doi:10.1172/jci.insight.99363.

30. M. Gonzalez-Gronow, S. V. Pizzo, Physiological Roles of the Autoantibodies to the 78-Kilodalton Glucose-Regulated Protein (GRP78) in Cancer and Autoimmune Diseases. Biomedicines 10, 1222 (2022).

31. H.-S. Hsu, C.-C. Liu, J.-H. Lin, T.-W. Hsu, J.-W. Hsu, K. Su, S.-C. Hung, Involvement of ER stress, PI3K/AKT activation, and lung fibroblast proliferation in bleomycin-induced pulmonary fibrosis. Sci. Rep. 7, 14272 (2017).

32. D. Vosoughi, A. Ulahannan, Q. Li, E. Huszti, A. Chruscinski, D. Birriel, G. Madu, G. Teskey, M. Aversa, T. Martinu, S. Juvet, Humoral immunity to lung antigens early post-transplant confers risk for chronic lung allograft dysfunction. J. Heart Lung Transplant. 0 (2025), doi:10.1016/j.healun.2025.02.1577.

33. P. Tondo, G. Scioscia, C. C. De Pace, F. Murgolo, F. Maci, G. M. Stella, D. Pescatore, M. P. Foschino Barbaro, D. Lacedonia, Gender Differences Are a Leading Factor in 5-Year Survival of Patients with Idiopathic Pulmonary Fibrosis over Antifibrotic Therapy Reduction. Life 15, 106 (2025).

34. T. Zaman, T. Moua, E. Vittinghoff, J. H. Ryu, H. R. Collard, J. S. Lee, Differences in Clinical Characteristics and Outcomes Between Men and Women With Idiopathic Pulmonary Fibrosis. Chest 158, 245–251 (2020).

35. Y. Xu, T. Mizuno, A. Sridharan, Y. Du, M. Guo, J. Tang, K. A. Wikenheiser-Brokamp, A.-K. T. Perl, V. A. Funari, J. J. Gokey, B. R. Stripp, J. A. Whitsett, Single-cell RNA sequencing identifies diverse roles of epithelial cells in idiopathic pulmonary fibrosis. JCI Insight 1 (2017), doi:10.1172/jci.insight.90558.

36. A. C. Habermann, A. J. Gutierrez, L. T. Bui, S. L. Yahn, N. I. Winters, C. L. Calvi, L. Peter, M.-I. Chung, C. J. Taylor, C. Jetter, L. Raju, J. Roberson, G. Ding, L. Wood, J. M. S. Sucre, B. W. Richmond, A. P. Serezani, W. J. McDonnell, S. B. Mallal, M. J. Bacchetta, J. E. Loyd, C. M. Shaver, L. B. Ware, R. Bremner, R. Walia, T. S. Blackwell, N. E. Banovich, J. A. Kropski, Single-cell RNA sequencing reveals profibrotic roles of distinct epithelial and mesenchymal lineages in pulmonary fibrosis. Sci. Adv. 6, eaba1972 (2020).

37. S. Y. Shin, J. Chen, I. Milman Krentsis, Y. Reisner, R. Abrencillo, R. Hussain, D. Wu, H. Karmouty-Quintana, From Epithelium to Therapy: Transitional Cells in Lung Fibrosis. Am. J. Respir. Cell Mol. Biol. 72, 472–483.

38. M. Strunz, L. M. Simon, M. Ansari, J. J. Kathiriya, I. Angelidis, C. H. Mayr, G. Tsidiridis, M. Lange, L. F. Mattner, M. Yee, P. Ogar, A. Sengupta, I. Kukhtevich, R. Schneider, Z. Zhao, C. Voss, T. Stoeger, J. H. L. Neumann, A. Hilgendorff, J. Behr, M. O’Reilly, M. Lehmann, G. Burgstaller, M. Königshoff, H. A. Chapman, F. J. Theis, H. B. Schiller, Alveolar regeneration through a Krt8+ transitional stem cell state that persists in human lung fibrosis. Nat. Commun. 11, 3559 (2020).

39. V. Bergen, M. Lange, S. Peidli, F. A. Wolf, F. J. Theis, Generalizing RNA velocity to transient cell states through dynamical modeling. Nat. Biotechnol. 38, 1408–1414 (2020).

40. P. Jiang, R. Gil de Rubio, S. M. Hrycaj, S. J. Gurczynski, K. A. Riemondy, B. B. Moore, M. B. Omary, K. M. Ridge, R. L. Zemans, Ineffectual Type 2–to–Type 1 Alveolar Epithelial Cell Differentiation in Idiopathic Pulmonary Fibrosis: Persistence of the KRT8hi Transitional State. Am. J. Respir. Crit. Care Med. 201, 1443–1447 (2020).

41. T. G. Coursey, J. Tukler Henriksson, F. L. Barbosa, C. S. de Paiva, S. C. Pflugfelder, Interferon-γ–Induced Unfolded Protein Response in Conjunctival Goblet Cells as a Cause of Mucin Deficiency in Sjögren Syndrome. Am. J. Pathol. 186, 1547–1558 (2016).

42. R. Choo-Wing, M. A. Syed, A. Harijith, B. Bowen, G. Pryhuber, C. Janér, S. Andersson, R. J. Homer, V. Bhandari, Hyperoxia and Interferon-γ–Induced Injury in Developing Lungs Occur via Cyclooxygenase-2 and the Endoplasmic Reticulum Stress–Dependent Pathway. Am. J. Respir. Cell Mol. Biol. 48, 749–757 (2013).

43. S.-A. Yoo, S. You, H.-J. Yoon, D.-H. Kim, H.-S. Kim, K. Lee, J. H. Ahn, D. Hwang, A. S. Lee, K.-J. Kim, Y.-J. Park, C.-S. Cho, W.-U. Kim, A novel pathogenic role of the ER chaperone GRP78/BiP in rheumatoid arthritis. J. Exp. Med. 209, 871–886 (2012).

44. P. A. Reyfman, J. M. Walter, N. Joshi, K. R. Anekalla, A. C. McQuattie-Pimentel, S. Chiu, R. Fernandez, M. Akbarpour, C.-I. Chen, Z. Ren, R. Verma, H. Abdala-Valencia, K. Nam, M. Chi, S. Han, F. J. Gonzalez-Gonzalez, S. Soberanes, S. Watanabe, K. J. N. Williams, A. S. Flozak, T. T. Nicholson, V. K. Morgan, D. R. Winter, M. Hinchcliff, C. L. Hrusch, R. D. Guzy, C. A. Bonham, A. I. Sperling, R. Bag, R. B. Hamanaka, G. M. Mutlu, A. V. Yeldandi, S. A. Marshall, A. Shilatifard, L. A. N. Amaral, H. Perlman, J. I. Sznajder, A. C. Argento, C. T. Gillespie, J. Dematte, M. Jain, B. D. Singer, K. M. Ridge, A. P. Lam, A. Bharat, S. M. Bhorade, C. J. Gottardi, G. R. S. Budinger, A. V. Misharin, Single-Cell Transcriptomic Analysis of Human Lung Provides Insights into the Pathobiology of Pulmonary Fibrosis. Am. J. Respir. Crit. Care Med. 199, 1517–1536 (2019).

45. N. Fujino, H. Kubo, T. Suzuki, C. Ota, A. E. Hegab, M. He, S. Suzuki, T. Suzuki, M. Yamada, T. Kondo, H. Kato, M. Yamaya, Isolation of alveolar epithelial type II progenitor cells from adult human lungs. Lab. Invest. 91, 363–378 (2011).

46. A. Duong, A. Wong, R. Ramendra, D. Sebben, S. Moshkelgosha, S. MacParland, M. Liu, S. Juvet, T. Martinu, A Rapid Human Lung Tissue Dissociation Protocol Maximizing Cell Yield and Minimizing Cellular Stress. Am. J. Respir. Cell Mol. Biol. 71, 509–518 (2024).

47. A. Murthy, L. R. Rodriguez, T. Dimopoulos, S. Bui, S. Iyer, K. Chavez, Y. Tomer, V. Abraham, C. Cooper, D. M. Renner, J. B. Katzen, I. D. Bentley, S. N. Ghadiali, J. A. Englert, S. R. Weiss, M. F. Beers, Activation of alveolar epithelial ER stress by β-coronavirus infection disrupts surfactant homeostasis in mice: implications for COVID-19 respiratory failure. Am. J. Physiol. -Lung Cell. Mol. Physiol. 327, L232–L249 (2024).

48. G. Saren, A. Wong, Y.-B. Lu, C. Baciu, W. Zhou, R. Zamel, S. Soltanieh, J. Sugihara, M. Liu, Ischemia-Reperfusion Injury in a Simulated Lung Transplant Setting Differentially Regulates Transcriptomic Profiles between Human Lung Endothelial and Epithelial Cells. Cells 10, 2713 (2021).

49. X.-Y. Zheng, H. Huang, Z.-T. Wei, H.-J. Yan, X.-W. Wang, L. Xu, C.-H. Li, H.-T. Tang, J.-J. Wang, Z.-W. Yu, D. Tian, Genetic effect of ischemia-reperfusion injury upon primary graft dysfunction and chronic lung allograft dysfunction in lung transplantation: evidence based on transcriptome data. Transpl. Immunol. 71, 101556 (2022).

50. R. N. Criner, E. Clausen, E. Cantu, Primary graft dysfunction. Curr. Opin. Organ Transplant. 26, 321–327 (2021).

51. T. Watanabe, T. Martinu, A. Chruscinski, K. Boonstra, B. Joe, M. Horie, Z. Guan, K. F. Bei, D. M. Hwang, M. Liu, S. Keshavjee, S. C. Juvet, A B cell-dependent pathway drives chronic lung allograft rejection after ischemia-reperfusion injury in mice. Am. J. Transplant. Off. J. Am. Soc. Transplant. Am. Soc. Transpl. Surg. 19, 3377–3389 (2019).

52. M. Keller, E. Bush, J. M. Diamond, P. Shah, J. Matthew, A. W. Brown, J. Sun, I. Timofte, H. Kong, I. Tunc, H. Luikart, A. Iacono, S. D. Nathan, K. K. Khush, J. Orens, M. Jang, S. Agbor-Enoh, Use of donor-derived-cell-free DNA as a marker of early allograft injury in primary graft dysfunction (PGD) to predict the risk of chronic lung allograft dysfunction (CLAD). J. Heart Lung Transplant. 40, 488–493 (2021).

53. T. Martinu, H. Oishi, S. C. Juvet, M. Cypel, M. Liu, G. J. Berry, D. M. Hwang, S. Keshavjee, Spectrum of chronic lung allograft pathology in a mouse minor-mismatched orthotopic lung transplant model. Am. J. Transplant. Off. J. Am. Soc. Transplant. Am. Soc. Transpl. Surg. 19, 247–258 (2019).

54. C. J. Iasella, A. Hoji, I. Popescu, J. Wei, M. E. Snyder, Y. Zhang, W. Xu, V. Iouchmanov, R. Koshy, M. Brown, M. Fung, C. Langelier, E. A. Lendermon, D. Dugger, R. Shah, J. Lee, B. Johnson, J. Golden, L. E. Leard, M. E. Kleinhenz, S. Kilaru, S. R. Hays, J. P. Singer, P. G. Sanchez, M. R. Morrell, J. M. Pilewski, J. R. Greenland, K. Chen, J. F. McDyer, Type-1 immunity and endogenous immune regulators predominate in the airway transcriptome during chronic lung allograft dysfunction. Am. J. Transplant. Off. J. Am. Soc. Transplant. Am. Soc. Transpl. Surg. 21, 2145–2160 (2021).

55. K. Moghbeli, I. Popescu, C. J. Iasella, S. Lieber, M. E. Snyder, M. Sciullo, R. Koshy, A. Zeng, R. P. Mohanty, C. Langelier, E. A. Lendermon, R. Shah, B. Johnson, D. Zhu, T. Martinu, L. E. Leard, M. E. Kleinhenz, S. R. Hays, J. P. Singer, N. Shigemura, J. M. Pilewski, C. A. Hage, K. Chen, J. R. Greenland, J. F. McDyer, Immune Gene Correlation Networks Differentiate Both Chronic Lung Allograft Dysfunction and Survival. Am. J. Transplant. 0 (2026), doi:10.1016/j.ajt.2026.01.025.

56. P. Xiao, Z. Hu, J. Lang, T. Pan, R. T. Mertens, H. Zhang, K. Guo, M. Shen, H. Cheng, X. Zhang, Q. Cao, Y. Ke, Mannose metabolism normalizes gut homeostasis by blocking the TNF-α-mediated proinflammatory circuit. Cell. Mol. Immunol. 20, 119–130 (2023).

57. A. M. Woodward, A. Di Zazzo, S. Bonini, P. Argüeso, Endoplasmic reticulum stress promotes inflammation-mediated proteolytic activity at the ocular surface. Sci. Rep. 10, 2216 (2020).

58. F. Shimizu, K. L. Schaller, G. P. Owens, A. C. Cotleur, D. Kellner, Y. Takeshita, B. Obermeier, T. J. Kryzer, Y. Sano, T. Kanda, V. A. Lennon, R. M. Ransohoff, J. L. Bennett, Glucose-regulated protein 78 autoantibody associates with blood-brain barrier disruption in neuromyelitis optica. Sci. Transl. Med. 9, eaai9111 (2017).

59. E. Murphy, C. Steenbergen, Mechanisms Underlying Acute Protection from Cardiac Ischemia-Reperfusion Injury. Physiol. Rev. 88, 581–609 (2008).

60. C. Li, Q. Ma, S. Toan, J. Wang, H. Zhou, J. Liang, SERCA overexpression reduces reperfusion-mediated cardiac microvascular damage through inhibition of the calcium/MCU/mPTP/necroptosis signaling pathways. Redox Biol. 36, 101659 (2020).

61. K. F. R. Pobre, G. J. Poet, L. M. Hendershot, The endoplasmic reticulum (ER) chaperone BiP is a master regulator of ER functions: Getting by with a little help from ERdj friends. J. Biol. Chem. 294, 2098–2108 (2019).

62. D. Ron, P. Walter, Signal integration in the endoplasmic reticulum unfolded protein response. Nat. Rev. Mol. Cell Biol. 8, 519–529 (2007).

63. G. Li, M. Mongillo, K.-T. Chin, H. Harding, D. Ron, A. R. Marks, I. Tabas, Role of ERO1-α– mediated stimulation of inositol 1,4,5-triphosphate receptor activity in endoplasmic reticulum stress–induced apoptosis. J. Cell Biol. 186, 783–792 (2009).

64. R. Rizzuto, D. De Stefani, A. Raffaello, C. Mammucari, Mitochondria as sensors and regulators of calcium signalling. Nat. Rev. Mol. Cell Biol. 13, 566–578 (2012).

65. E. T. Chouchani, V. R. Pell, E. Gaude, D. Aksentijević, S. Y. Sundier, E. L. Robb, A. Logan, S. M. Nadtochiy, E. N. J. Ord, A. C. Smith, F. Eyassu, R. Shirley, C.-H. Hu, A. J. Dare, A. M. James, S. Rogatti, R. C. Hartley, S. Eaton, A. S. H. Costa, P. S. Brookes, S. M. Davidson, M. R. Duchen, K. Saeb-Parsy, M. J. Shattock, A. J. Robinson, L. M. Work, C. Frezza, T. Krieg, M. P. Murphy, Ischaemic accumulation of succinate controls reperfusion injury through mitochondrial ROS. Nature 515, 431–435 (2014).

66. J. M. McCord, Oxygen-derived free radicals in postischemic tissue injury. N. Engl. J. Med. 312, 159–163 (1985).

67. H. K. Eltzschig, T. Eckle, Ischemia and reperfusion--from mechanism to translation. Nat. Med. 17, 1391–1401 (2011).

68. H. Dridi, G. Santulli, L. Bahlouli, M. C. Miotto, G. Weninger, A. R. Marks, Mitochondrial Calcium Overload Plays a Causal Role in Oxidative Stress in the Failing Heart. Biomolecules 13, 1409 (2023).

69. H. Kim, J. Zhao, Q. Zhang, Y. Wang, D. Lee, X. Bai, L. Turrell, M. Chen, W. Gao, S. Keshavjee, M. Liu, δV1-1 Reduces Pulmonary Ischemia Reperfusion-Induced Lung Injury by Inhibiting Necrosis and Mitochondrial Localization of PKCδ and p53. Am. J. Transplant. Off. J. Am. Soc. Transplant. Am. Soc. Transpl. Surg. 16, 83–98 (2016).

70. S. S. Cao, R. J. Kaufman, Endoplasmic Reticulum Stress and Oxidative Stress in Cell Fate Decision and Human Disease. Antioxid. Redox Signal. 21, 396–413 (2014).

71. A. L. Eiliazadeh, M. A. V. Suma, J. Li, M. Cuesta, J. Ishaq, D. Taniguchi, S. Keshavjee, T. K. Waddell, G. Karoubi, High-throughput isolation of normal and pulmonary fibrosis human alveolar epithelial cells. Sci. Rep. 16, 757 (2025).

72. L. Levy, S. C. Juvet, K. Boonstra, L. G. Singer, S. Azad, B. Joe, M. Cypel, S. Keshavjee, T. Martinu, Sequential broncho-alveolar lavages reflect distinct pulmonary compartments: clinical and research implications in lung transplantation. Respir. Res. 19, 102 (2018).

73. T. Watanabe, S. C. Juvet, G. Berra, J. Havlin, W. Zhong, K. Boonstra, T. Daigneault, M. Horie, C. Konoeda, G. Teskey, Z. Guan, D. M. Hwang, M. Liu, S. Keshavjee, T. Martinu, Donor IL-17 receptor A regulates LPS-potentiated acute and chronic murine lung allograft rejection. JCI Insight 8, e158002.

74. S. Moshkelgosha, L. Levy, S. Safavi, S. Karunagaran, G. Wilson, B. Renaud-Picard, G. Madu, R. Ramchandani, J. Oliver, T. Watanabe, K. F. Bei, B. Joe, Q. Li, E. Huszti, M. Cheung, D. Hedley, J. Yeung, S. Keshavjee, T. Martinu, S. Juvet, Emergence of a senescent and inflammatory pulmonary CD4+ T cell population prior to lung allograft failure. Sci. Adv. 11, eadp9052 (2025).

75. T. Watanabe, C. Lam, J. Oliver, H. Oishi, G. Teskey, S. Beber, K. Boonstra, J. Mauricio Umaña, H. Buhari, B. Joe, Z. Guan, M. Horie, S. Keshavjee, T. Martinu, S. C. Juvet, Donor Batf3 inhibits murine lung allograft rejection and airway fibrosis. Mucosal Immunol. 16, 104– 120 (2023).

