## Supplementary Materials and Methods for "The unfolded protein response in alveolar type II cells is a therapeutic target in ischemia-reperfusion injury and chronic lung allograft rejection"

**Supplemental Table 1.** List of antigens used in this study.

| **Antigen Name (Used in Assay)** | **Full name** | **Explanation / Species** | **Company** | **Company Location** | **Catalog ID** |
| --- | --- | --- | --- | --- | --- |
| A1AR | Alpha 1A adrenergic receptor (A1AR) fragment | Peptide | GenScript | Piscataway, NJ 08854 | |
| Actin (Bovine) | Actin | Bovine muscle | Sigma | Saint Louis, MO 63103 | A3653 |
| Actin Rabbit | Actin (rabbit muscle) | Rabbit muscle | Molecular Probes | Waltham, MA USA 02451 | |
| Aggrecan | Aggrecan | Bovine articular cartilage | Sigma | Saint Louis, MO 63103 | A1960 |
| Aldolase, Type X | Aldolase, Type X | Rabbit muscle | Sigma | Saint Louis, MO 63103 | A2714 |
| Alpha elastin | Alpha-elastin | Recombinant, human | Dr. Keeley (University of Toronto) | Toronto, Ontario M5S 3M2 | |
| alpha-2-macroglobulin | Recombinant Human Alpha-2-Macroglobulin/A2M Protein | Human | RayBiotech | Peachtree Corners, GA, 30092 | 230-30115-50 |
| alpha-Actinin | Alpha-actinin | Chicken gizzard | Sigma | Saint Louis, MO 63103 | A9776 |
| Alpha-KGDH | Alpha-ketoglutarate dehydrogenase | Porcine heart | Sigma | Saint Louis, MO 63103 | K1502 |
| alphaB-crystallin | Alpha B-crystallin | Recombinant, human | Enzo (stressgen) | San Diego, California 92121 | |
| AQP1 | Aquaporin-1 (AQP1) Recombinant Protein | Human | MyBioSource | San Diego, CA 92195-3308 | MBS948542 |
| AT1 | angiotensin 2 receptor (AT1R) fragment | Peptide | GenScript | Piscataway, NJ 08854 | |
| B1AR | Beta 1 adrenergic receptor (B1AR) fragment | Peptide | GenScript | Piscataway, NJ 08854 | |
| B2AR | Beta 2 adrenergic receptor (B2AR) fragment | Peptide | GenScript | Piscataway, NJ 08854 | |
| bgal | β-galactosidase | | Creative Enzymes | |  |
| BPI | Bactericidal/Permeability Increasing protein | Non-recombinant, human | Diarect | Eden Prairie, MN 55344 | 19200 |
| C1q (purified non-recoinant) | Complement component C1q | Purified from human serum | Sigma | Saint Louis, MO 63103 | C1740 |
| Cardiolipin C1649 | Cardiolipin | Bovine heart, solution in ethanol | Sigma | Saint Louis, MO 63103 | C1649 |
| CENP-A | Centromere Protein A | Recombinant, human | Diarect | Eden Prairie, MN 55344 | 16900 |
| CENP-B | Centromere Protein B | Recombinant, human | Diarect | Eden Prairie, MN 55344 | 12500 |
| CLDN5 | Recombinant Human CLDN5 | Human | Biorbyrt | Cambridge, United Kingdom, CB5 8LA | orb1099642 |
| CLU | Human Clusterin (CLU) Protein | Human | Abbexa | Cambridge, United Kingdom, CB4 0GJ | abx065948 |
| Collagen I | Collagen, I | Human placenta | Sigma | Saint Louis, MO 63103 | C7774 |
| Collagen III | Collagen, III | Human placenta | Sigma | Saint Louis, MO 63103 | C4407 |
| Collagen IV | Collagen, IV | Human placenta | Sigma | Saint Louis, MO 63103 | C5533 |
| Collagen V | Collagen, V | Human placenta | Sigma | Saint Louis, MO 63103 | C3657 |
| Collagen VI | Collagen, VI | Human placenta | Sigma | Saint Louis, MO 63103 | C7521 |
| Desmin | Desmin | Recombinant, human | GenWay | San Diego, CA 92121, United States | GWB-2E8E72 |
| dsDNA (genomic) | Deoxyribonucleic acid, double stranded | Salmon testes, sodium salt | Sigma | Saint Louis, MO 63103 | D1626 |
| dsDNA (plasmid) | DNA circualar plasmid, Double Stranded | Bacterial plasmid | Diarect | Eden Prairie, MN 55344 | 12300 |
| Ebna peptide | EBV nuclear antigen | 398-412 peptide | Stanford | Stanford, CA |  |
| EGFR | Recombinant Human EGFR/ERBB1/HER1 | Human | RayBiotech | Peachtree Corners, GA, 30092 | 230-30016-50 |
| FBN1 | Human Fibrillin 1 (FBN1) Protein | Human | Abbexa | Cambridge, United Kingdom, CB4 0GJ | abx066564 |
| Fib I | Fibrinogen, type I | Human plasma | Sigma | Saint Louis, MO 63103 | F3879 |
| Fib I-S | Fibrinogen, type I-S | Bovine plasma | Sigma | Saint Louis, MO 63103 | F8630 |
| GBM, diss | Glomerular Basement Membrane (GBM; dissociated) | Recombinant, human | Diarect | Eden Prairie, MN 55344 | 16800 |
| GP2 | Zymogen granule membrane glycoprotein 2 | Recombinant, human | Diarect | Eden Prairie, MN 55344 | 19600 |
| gp210 | glycoprotein 210 | Recombinant, human | Diarect | Eden Prairie, MN 55344 | 19000 |
| Grp78/BiP | GRP78 (HSPa5 or BiP) immunoglobulin heavy chain-binding protein | Recombinant hamster | Enzo (Stressgen) | San Diego, California 92121 | ADI-SPP-765 |
| HCEC cytop | Human Cardiac Endothelial Cells Lysate | Human |  |  |  |
| HCEC memb | Human Cardiac Endothelial Cell Lysate | Human |  |  |  |
| HCEC total (SDS) | Human Cardiac Endothelial Cell Lysate | Human |  |  |  |
| HCEC total (Triton) | Human Cardiac Endothelial Cell Lysate | Human |  |  |  |
| Heparin | Heparin Sulfate | Bovine kidney, sodium salt | Sigma | Saint Louis, MO 63103 | H7640 |
| Histone H2A-H2B dimers | Histone H2A-H2B dimers | Recombinant, human | EMDMillipore | Saint Louis, MO 63103 | 14-1052 |
| Histone H2B | Histone H2B | Recombinant, human | Active Motif | Carlsbad, CA 92008 | 31492 |
| Histone H3 (1-136 aa) | Histone H3 (1-136 aa) | Recombinant, human | RayBiotech | Peachtree Corners, GA 30092 | 268-11222-1 |
| Histone H4 (1-103 aa) | Histone H4 (1-103 aa) | Recombinant, human | RayBiotech | Peachtree Corners, GA 30092 | 268-11223-1 |
| HMG CoA | 3-hydroxy-3-methylglutaryl-Coenzyme A Reductase | Recombinant, human | Cayman | Ann Arbor, Michigan 48108 | 14944 |
| HPAEpiC | Human Pulmonary Alveolar Epithelial Cell Lysate | Human | ScienCell Research Laboratories | Carlsbad, CA 92008 | 3206 |
| HPMEC | Human Pulmonary Microvascular Endothelial Cell Lysate | Human | ScienCell Research Laboratories | Carlsbad, CA 92008 | 3006 |
| HSAEpiC | Human Pulmonary Small Airway Epithelial Cell Lysate | Human | ScienCell Research Laboratories | Carlsbad, CA 92008 | 3236 |
| HSP 47 | Heat shock protein 47 | Recombinant, human | Enzo (Stressgen) | San Diego, California 92121 | ADI-SPP-535 |
| HSP 60 | Heat shock protein 60 | Recombinant, human | Enzo (Stressgen) | San Diego, California 92121 | ADI-NSP-540-E |
| HSP 90 | Heat shock protein 90 | Native Human, from HeLa Cells | Enzo (Stressgen) | San Diego, California 92121 | ADI-SPP-770 |
| human albumin | Human albumin | recombinant, expressed in rice, lyophilized powder, suitable for cell culture, low endotoxin, ≥96% (SDS-PAGE) | Sigma | Saint Louis, MO 63103 | A9731 |
| human C1q (Abcam) | Native Human C1q protein | Source: plasma | Abcam | Cambridge, CB2 0AX, UK | ab96363 |
| Human core histones | Human core histones | Purified from Hela cells | RayBiotech | Peachtree Corners, GA 30092 | 268-11229-1 |
| human elastin | Recombinant, human elastin | Human | Dr. Keeley (University of Toronto) | Toronto, Ontario M5S 3M2 | |
| human fgl2 | fibrinogen-like 2 | Recombinant, human | Dr. Gary Levy (University Health Network) | Toronto, ON M5G 2C4 | |
| Human H1 Chromatin | Human H1 Chromatin | | Dr. Joan Wither(University Health Network) | Toronto, ON M5G 2C4 | |
| Human IgA | Human IgA | Purified from human serum | Jackson Immunoresearch | West Grove, PA 19390 | 009-000-011 |
| Human IgE | Human IgE | From a monoclonal hybridoma, human | Abcam | Cambridge, CB2 0AX, UK | ab65866 |
| Human IgG | Human IgG | ChromPure human IgG, whole molecule | Jackson Immunoresearch | West Grove, PA 19390 | 009-000-003 |
| Human IgM | Human IgM | ChromPure human IgM, myeloma | Jackson Immunoresearch | West Grove, PA 19390 | 009-000-012 |
| human intestinal smooth muscle cell | Human Intestinal Smooth Muscle Cell Lysate | Lysate from primary human cells | ScienCell Research Laboratories | Carlsbad, CA 92008 | |
| human nucleosome | Human nucleosome | Purified from Hela cells | EpiGex | 67412 Illkirch cedex France | EPX-01-NCN |
| IGFBP4 | IGFBP4 Protein, Human, Recombinant (His Tag) | Human | Sino Biological | China | 10967-H08H |
| Insulin | Insulin | Recombinant, human | Sigma | Saint Louis, MO 63103 | I2643 |
| Jo-1 | Histidyl-tRNA synthetase | Recombinant, human | Diarect | Eden Prairie, MN 55344 | 12900 |
| Ku (p70/p80) | Ku (p70/p80) | Recombinant, human | Diarect | Eden Prairie, MN 55344 | 17300 |
| La (SS-B) Antigens (Immunovision) | La (SS-B) Antigens (47 kD ribonucleic protein) | Purified from calf and/or rabbit thymus | Immunovision | Springdale, Arkansas 72764 | SSB-3000 |
| La/SS-B (Recombinant, human diarect) | La (SS-B) Antigens (47 kD ribonucleic protein) | Recombinant, human | Diarect | Eden Prairie, MN 55344 | 12800 |
| Laminin | Laminin | Engelbreth-Holm-Swarm murine sarcoma basement membrane | Sigma | Saint Louis, MO 63103 | L2020 |
| LCI | liver cytosol type 1 antigen | Recombinant, human | Diarect | Eden Prairie, MN 55344 | 13700 |
| LG3 | immunogenic fragment of perlecan | Recombinant, human | Dr. Marie-Josee Hebert (University of Montreal) | Montreal, Quebec H3T 1J4 | |
| LPS | Lipopolysaccharides from Escherichia coli O111:B4 | E.Coli | Sigma | St. Louis, MO 63103 | L2630 |
| M2 | The branched-chain α-ketoacid dehydrogenase complex: BCKDC or BCKDH complex) | Recombinant, human | Diarect | Eden Prairie, MN 55344 | 18000 |
| Measles | Measles antigen | Edmonston Strain ATCC #VR-24 | Meridian | The Woodlands, TX 77380 | R14120 |
| Mi-2 | Subunit of the nucleosome remodeling deacetylase (NuRD) complex | Recombinant, human | Diarect | Eden Prairie, MN 55344 | 18100 |
| mouse fgl2 | Mouse fibrinogen-like 2 | Recombinant, mouse | Dr. Gary Levy (University Health Network) | Toronto, ON M5G 2C4 | |
| MP (Antigona) | 2-Methacryloyloxyethyl phosphorylcholine | | Sigma | Saint Louis, MO 63103 | 730114-5G |
| MPO | Myeloperoxidase (MPO) | Nonrecombinant, human | Diarect | Eden Prairie, MN 55344 | 18500 |
| myh6 | Recombinant Human Myosin-6 (MYH6), partial | Human | CUSABIO | Houston, TX 77054 | CSB-YP015299HU |
| Myosin | Myosin, Calcium activated from porcine heart | Porcine | Sigma | St. Louis, MO 63103 | M0531 |
| Myosin Bind Protein C | Myosin Binding Protein C | Recombinant, human | Dr. Sadayappan (Loyola University) | Chicago, IL 60660 | |
| NAPSA | NAPSA Recombinant Protein (Human) | Human | Aviva Systems Biology Corporation | San Diego, CA 92121 | OPCA01745 |
| NPNT | Human Nephronectin (NPNT) Protein | Human | Abbexa | Cambridge, United Kingdom, CB4 0GJ | abx068217 |
| Nucleolin | Nucleolin | Recombinant, human | Diarect | Eden Prairie, MN 55344 | 19700 |
| OGDC-E2 | 2-oxoglutarate dehydrogenase complex | Recombinant, human | Diarect | Eden Prairie, MN 55344 | 17800 |
| Ox LDL | Copper Oxidized human LDL | Human plasma | Academy Bio-Medical Company | Houston, TX 77020 | 20P-OX-L102 |
| PBS | phosphate buffered saline | | Wisent Bioproducts | Saint-Jean-Baptiste, QC J0L 2B0 | |
| PCNA | Proliferating cell nuclear antigen | Recombinant, human | Diarect | Eden Prairie, MN 55344 | 36600 |
| PDC-E2 | E2 component of mitochondrial pyruvate dehydrogenase complex | Recombinant, human | Diarect | Eden Prairie, MN 55344 | |
| PDH | Pyruvate dehydrogenase | Porcine heart | Sigma | Saint Louis, MO 63103 | P7032 |
| PL-12 | Alanyl-tRNA Synthase (PL-12) | Recombinant, human | Diarect | Eden Prairie, MN 55344 | 15700 |
| PL-7 | Threonyl-tRNA Synthase (PL-7) | Recombinant, human | Diarect | Eden Prairie, MN 55344 | 15600 |
| PM/Scl 100 | polymyositis/scleroderma (PM/Scl) complex 100 | Recombinant, human | Diarect | Eden Prairie, MN 55344 | 16000 |
| PM/Scl 75 | polymyositis/scleroderma (PM/Scl) complex 75 | Recombinant, human | Diarect | Eden Prairie, MN 55344 | 17000 |
| PR3 | Proteinase 3 | Non-recomb human leukocytes | Diarect | Eden Prairie, MN 55344 | PR3 |
| Proteoglycan | Proteoglycan | Bovine nasal septum | Sigma | Saint Louis, MO 63103 | P5864 |
| PTC (Antigona) | L-α-Phosphatidylcholine | egg yolk, Type XVI-E, ≥99% (TLC), lyophilized powder | Sigma | Saint Louis, MO 63103 | L-α-P3556-25MG |
| Recombinant Histone H2A (hu) | Recombinant Histone H2A (hu) | Recombinant, human | Active Motif | Carlsbad, CA 92008 | 31490 |
| Ribo P0 | Ribosomal Phosphoprotein P0 | Recombinant, human | Diarect | Eden Prairie, MN 55344 | 14100 |
| RNP/Sm (non-recombinant; bovine) | RNP/Sm (non-recombinant; bovine) | Non-recombinant, bovine | Diarect | Eden Prairie, MN 55344 | 11600 |
| Ro/SS-A (52 kDa, human recombinant) | Ro/SS-A (52 kDa) | Recombinant, human | Diarect | Eden Prairie, MN 55344 | 12700 |
| Ro/SS-A (60 kD; non-recombinant; bovine) | Ro/SS-A (60 kDa) | Non-recombinant bovine | Diarect | Eden Prairie, MN 55344 | 15500 |
| Ro/SS-A (60kD; recombinant) | Ro/SS-A (60 kDa) | Recombinant, human | Diarect | Eden Prairie, MN 55344 | 17400 |
| Sars-CoV-2 RBD | SARS-CoV-2 (COVID-19, 2019-nCoV) Spike RBD Recombinant Protein | Human | ProSci Inc | Poway, CA 92064 | 10-100 |
| Sars-CoV2 Spike | SARS-CoV-2 (COVID-19, 2019-nCoV) Spike-RBD Recombinant Protein | Human | ProSci Inc | Poway, CA 92064 | 10-015 |
| SCGB1A1 | Secretoglobin family 1A member 1 | Recombinant, human | NBP2-35126) | exclude maybe | NBP2-35126 |
| Scl-70: trunc | DNA Topoisomerase I (Scl-70) truncated | Recombinant, human | Diarect | Eden Prairie, MN 55344 | 14500 |
| SFTPA1 | SFTPA1 protein :: Pulmonary Surfactant Associated Protein A1 (SFTPA1) Protein | Human | MyBioSource | San Diego, CA 92195-3308 | MBS2124300 |
| SLC34A2 | Human SLC34A2 protein | Human | Biorbyrt | Cambridge, United Kingdom, CB5 8LA | orb419212 |
| SmD | SmD | Recombinant, human | Diarect | Eden Prairie, MN 55344 | 11700 |
| SmD1 | Human small nuclear ribonucleoprotein D1 polypeptide (SNRPD1; SmD1). | Recombinant, human | Diarect | Eden Prairie, MN 55344 | 11800 |
| SmD2 | Human small nuclear ribonucleoprotein D2 polypeptide (SNRPD2; SmD2). | Recombinant, human | Diarect | Eden Prairie, MN 55344 | 11900 |
| SmD3 | Human small nuclear ribonucleoprotein D3 polypeptide (SNRPD3; SmD3). | Recombinant, human | Diarect | Eden Prairie, MN 55344 | 12000 |
| snRNP 68 B/B | U1-snRNP 68 Protein B/B | Recombinant, human | Diarect | Eden Prairie, MN 55344 | 13300 |
| snRNP A | U1-snRNP A Protein | Recombinant, human | Diarect | Eden Prairie, MN 55344 | 13100 |
| snRNP C | U1-snRNP C Protein | Recombinant, human | Diarect | Eden Prairie, MN 55344 | 13200 |
| SP-D | Surfactant protein D | Recombinant, human | Novusbio | Centennial, CO 80112 | 1920-SP |
| Sp100 | Sp100 nuclear antigen (Speckled 100 kDa) | Recombinant, human | Diarect | Eden Prairie, MN 55344 | 18900 |
| SRP54 | signal recognition particle (SRP) 54 GTPase protein | Recombinant, human | Diarect | Eden Prairie, MN 55344 | 18400 |
| ssDNA | Deoxyribonucleic acid, single stranded | Calf thymus | Sigma | Saint Louis, MO 63103 | D8899 |
| Testis Lysate | Human Testis Protein (Cell/Tissue Lysate) | Human | Lifespan Biosciences | Newark, CA | LS-G134108-0.1 |
| TGFBR2 | TGFBR2 Protein, Human, Recombinant (ECD, His Tag) | Human | Sino Biological | China | 10358-H08B |
| thyroglobulin #2 | dimeric glycoprotein thyroglobulin | Native. Isolated from human thyroid glands. | Diarect | Eden Prairie, MN 55344 | A12200 |
| Thyrogobulin | dimeric glycoprotein thyroglobulin | Native. Isolated from human thyroid glands. | Diarect | Eden Prairie, MN 55344 | A12200 |
| TPO | Thyroid peroxidase (TPO) | Recombinant, human | Diarect | Eden Prairie, MN 55344 | 12100 |
| Tropoelastin | Tropoelastin | Tropoelastin human recombinant, expressed in E. coli | Sigma | Saint Louis, MO 63103 | T0706 |
| Tropomyosin | Tropomyosin | Porcine muscle | Sigma | Saint Louis, MO 63103 | T2400 |
| Troponin C | Troponin C | Recombinant, human | CalBioreagents | San Mateo, CA 94403 | A093 |
| Troponin T | Troponin T | Human heart | Sigma | Saint Louis, MO 63103 | T0175 |
| tTG baculovirus | tTG (Tissue transglutaminase) baculovirus | Recombinant, human | Diarect | Eden Prairie, MN 55344 | 15200 |
| tuba1a | Recombinant Human TUBA1A | Human | RayBiotech | Peachtree Corners, GA, 30092 | 230-00849-50 |
| TUBA1B protein | Tubulin alpha-1B chain | Recombinant, human | Abnova | Taoyuan City 320 Taiwan | H00010376-P01 |
| Vimentin | Vimentin | Recombinant, human | Cedarlane | Burlington, ON L7L 5R2 | CLPR0309 |
| whole histones | Whole Histones | Purified from chicken RBC | Immunovision | Springdale, Arkansas 72764 | HIS-1000 |
| Mouse IgG | ChromPure® Mouse IgG, whole molecule | Jackson Immunoresearch | Highly purified serum proteins, non-immunized mice | West Grove, PA 19390 | 015-000-003 |
| Mouse GRP78 | Recombinant Mouse HSPA5/BIP Protein | Recombinant, mouse | MyBioSource | San Diego, CA 92195-3308 | MBS1578595 |

**Supplemental Table 2.** Characteristics of the nested case-control subset and Time-to-CLAD cohort.

|  | | **LT-FDC Cohort** | **Longitudinal Cohort** |
| --- | --- | --- | --- |
| Patient Count (n) | | 159 | 79 |
| Recipient Age at LT^a^ | | 52.1 [46.5, 61] | 54.2 [31.5, 58.5] |
| Donor Age at LT^a^ | | 46.4 [31, 60.5] | 45.2 [31.5, 58.5] |
| Donor Sex = M | | 89.0 (56.0) | 42.0 (53.2) |
| Recipient Sex = M | | 88.0 (55.3) | 45.0 (57.0) |
| Primary Disease | |  |  |
|  | Cystic Fibrosis | 23.0 (14.5) | 10.0 (12.7) |
|  | Emphysema/COPD | 46.0 (28.9) | 22.0 (27.8) |
|  | Pulmonary Fibrosis | 55.0 (34.6) | 35.0 (44.3) |
|  | Other | 35.0 (22.0) | 12.0 (15.2) |
| CMV Primary Mismatch | |  |  |
|  | R-/D- | 37.0 (23.3) | 29.0 (36.7) |
|  | R-/D+ | 32.0 (20.1) | 8.0 (10.1) |
|  | R+ (R+/D- or R+/D+) | 90.0 (56.6) | 42.0 (53.2) |
| Status at Tx Admission^b^ | |  |  |
|  | 1 | 38.0 (23.9) | 29.0 (36.7) |
|  | 2 | 68.0 (42.8) | 8.0 (10.1) |
|  | 3 | 52.0 (32.7) | 42.0 (53.2) |
| Follow Up Days^a^ | | 2266.3 [1014, 3524] | 2151.4 [1280.5, 2950.5] |

Values expressed as number (%) except where indicated.

^a^Values expressed as median [interquartile range].

^b^Where 1 reflects lowest acuity, 2 reflects moderate acuity, and 3 reflects highest acuity.

COPD, chronic obstructive pulmonary disease; CMV, cytomegalovirus; LT, lung transplant; Tx, transplant; FDC, feature discovery cohort.

**Supplemental Table 3.** Differential gene expression comparing AT2 Active and AT2 Homeostatic cell clusters in donor lungs.

|  | **Cluster** | **Gene** | **Average log2FC** | **PCT.1** | **PCT.2** | ***p*-Value Adjusted** |
| --- | --- | --- | --- | --- | --- | --- |
| **SFTPC** | AT2 Active | *SFTPC* | 0.486 | 0.998 | 0.999 | 0E+00 |
| **SLPI** | AT2 Active | *SLPI* | 0.563 | 1 | 1 | 3E-234 |
| **NDUFA4** | AT2 Active | *NDUFA4* | 0.347 | 0.954 | 0.986 | 2E-118 |
| **FABP5** | AT2 Active | *FABP5* | 0.542 | 0.987 | 0.996 | 2E-101 |
| **SCGB3A1** | AT2 Active | *SCGB3A1* | 0.741 | 0.738 | 0.63 | 8E-92 |
| **COX6A1** | AT2 Active | *COX6A1* | 0.366 | 0.902 | 0.951 | 9E-85 |
| **PLA2G1B** | AT2 Active | *PLA2G1B* | 0.600 | 0.826 | 0.842 | 2E-84 |
| **LRRC75A** | AT2 Active | *LRRC75A* | 0.906 | 0.576 | 0.488 | 5E-79 |
| **FOS** | AT2 Active | *FOS* | 0.459 | 0.905 | 0.915 | 2E-66 |
| **NDUFA13** | AT2 Active | *NDUFA13* | 0.334 | 0.852 | 0.922 | 2E-53 |
| **TTN** | AT2 Homeostatic | *TTN* | -0.922 | 0.815 | 0.964 | 0E+00 |
| **MALAT1** | AT2 Homeostatic | *MALAT1* | -0.398 | 0.995 | 1 | 2E-277 |
| **NEAT1** | AT2 Homeostatic | *NEAT1* | -0.569 | 0.988 | 1 | 7E-274 |
| **MACF1** | AT2 Homeostatic | *MACF1* | -0.693 | 0.647 | 0.887 | 2E-249 |
| **ARFGEF3** | AT2 Homeostatic | *ARFGEF3* | -0.684 | 0.589 | 0.847 | 2E-221 |
| **CCDC141** | AT2 Homeostatic | *CCDC141* | -0.762 | 0.538 | 0.805 | 2E-216 |
| **TANC2** | AT2 Homeostatic | *TANC2* | -0.711 | 0.495 | 0.78 | 7E-214 |
| **LRRFIP1** | AT2 Homeostatic | *LRRFIP1* | -0.503 | 0.829 | 0.964 | 9E-214 |
| **STEAP4** | AT2 Homeostatic | *STEAP4* | -0.747 | 0.662 | 0.885 | 1E-211 |
| **RNF145** | AT2 Homeostatic | *RNF145* | -0.492 | 0.891 | 0.982 | 2E-206 |

Differential gene expression between AT2 Active and AT2 Homeostatic clusters in DL samples were compared using the Wilcoxon rank-sum test in Seurat. Top genes reflect a surfactant-associated secretory program in AT2 Active cells and a structural gene expression program in AT2 Homeostatic cells. FC, fold-change; PCT, percent of cells expressing the gene in each group (PCT.1, AT2 Active; PCT.2, AT2 Homeostatic); *p*-values were adjusted using the Bonferroni correction method.

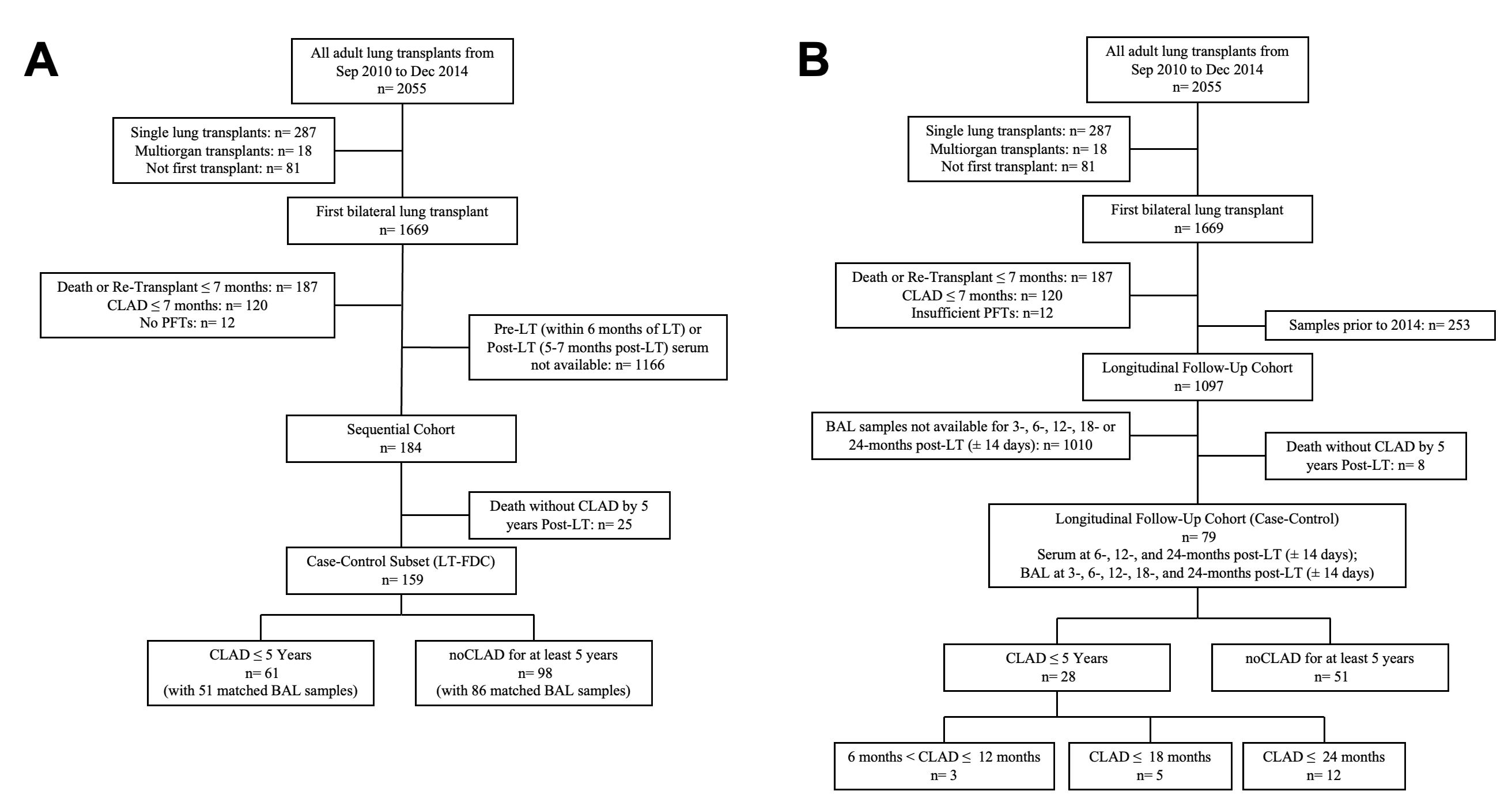

**Supplemental Figure 1. CONSORT diagrams depicting patient selection and cohort structure.** **A)** Flow diagram of inclusion and exclusion criteria for the LT-Feature Discovery Cohort (LT-FDC), with patients classified as having developed CLAD within 5 years post-transplant or remained CLAD-free for at least 5 years with sufficient follow-up. **B)** Overview of the longitudinal cohort design and BAL sample availability, applying the same case-control classification and indicating the number of patients who developed CLAD prior to each timepoint of interest.

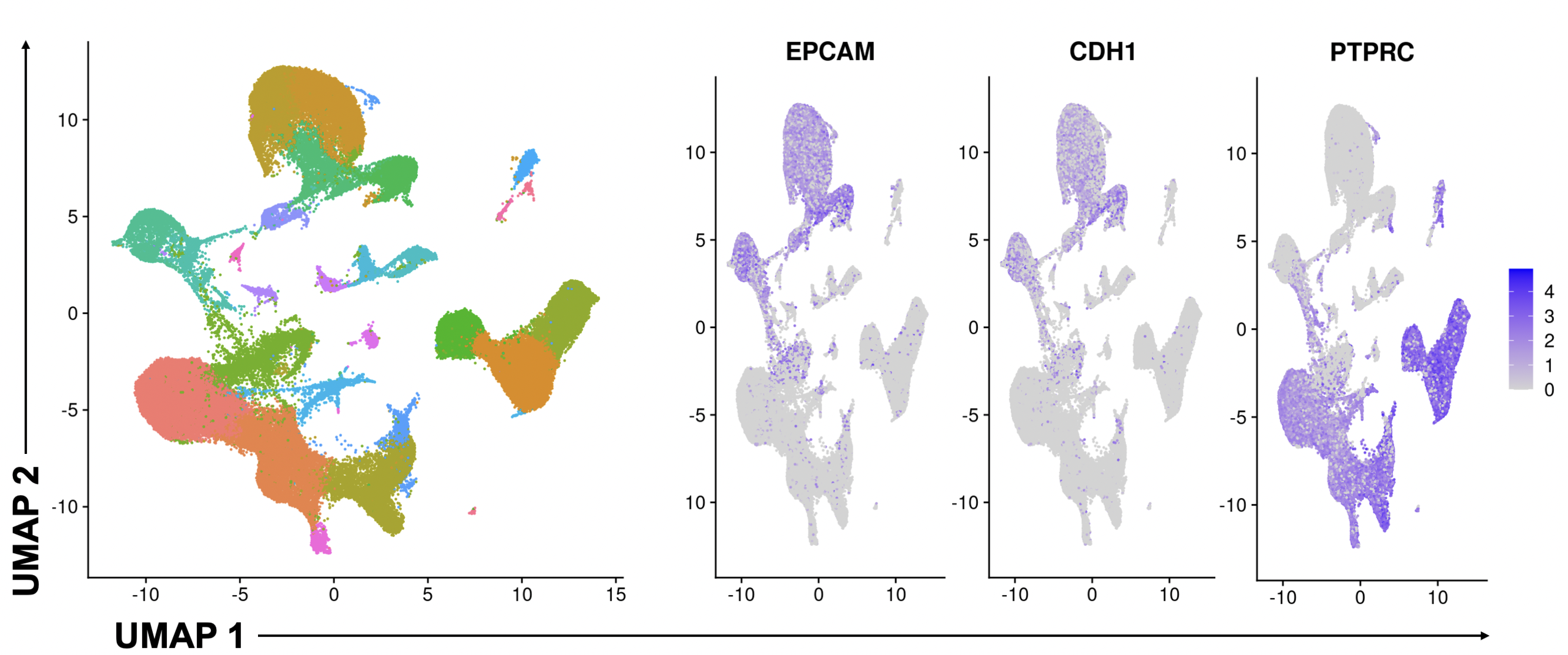

**Supplemental Figure 2. Epithelial cell isolation from scRNA-seq data.** Clusters were subsetted from integrated DL and CLAD scRNA-seq data sets based on canonical epithelial markers (*EPCAM, CDH1*), with *PTPRC* used to exclude hematopoietic lineage cells.

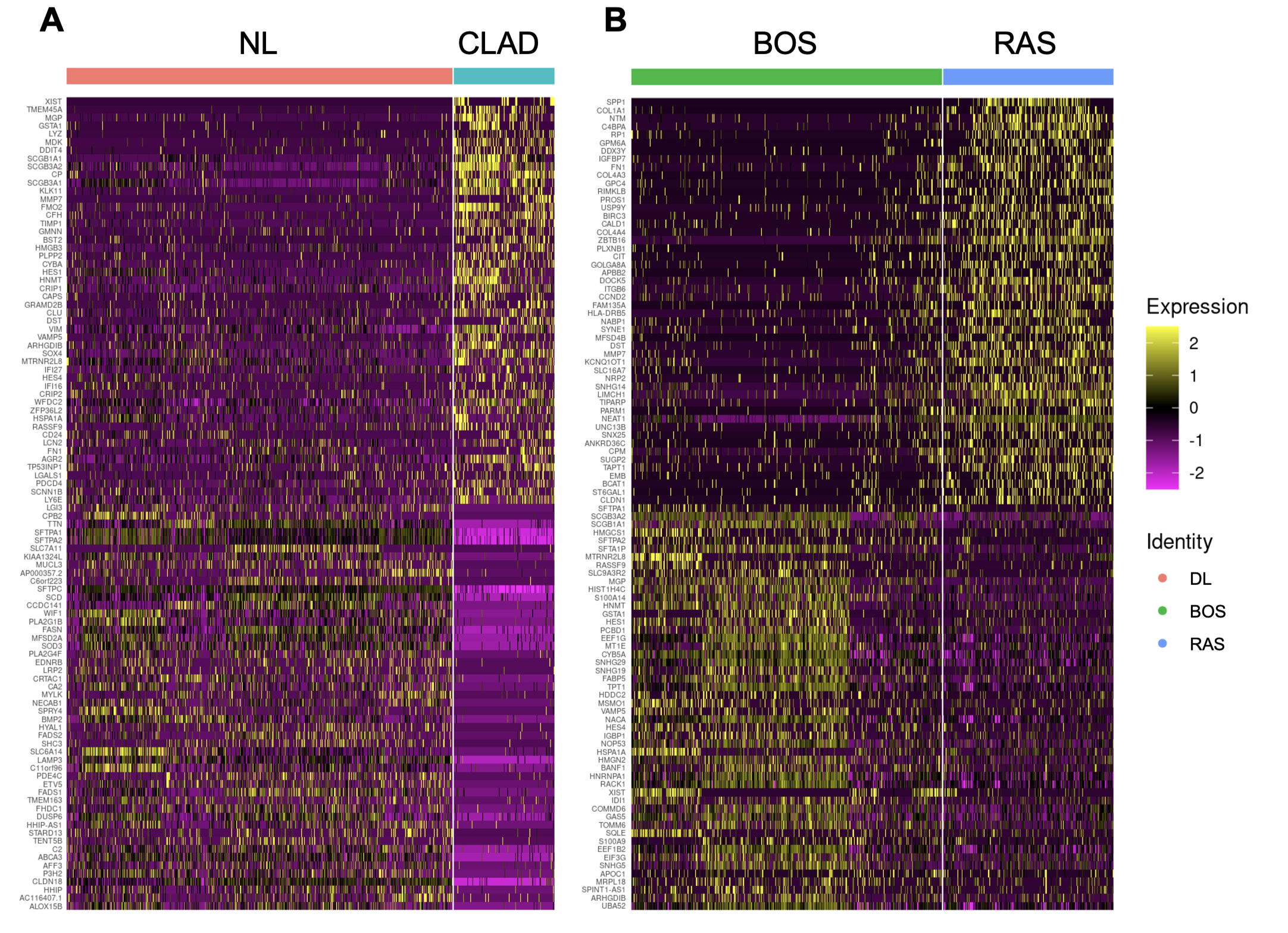

**Supplemental Figure 3. Differential gene expression in transitional AT2 cells.** **A)** Heatmap of the top 100 differentially expressed genes in transitional AT2 cells comparing CLAD and DL samples. **B)** Heatmap of the top 100 differentially expressed genes comparing BOS and RAS.

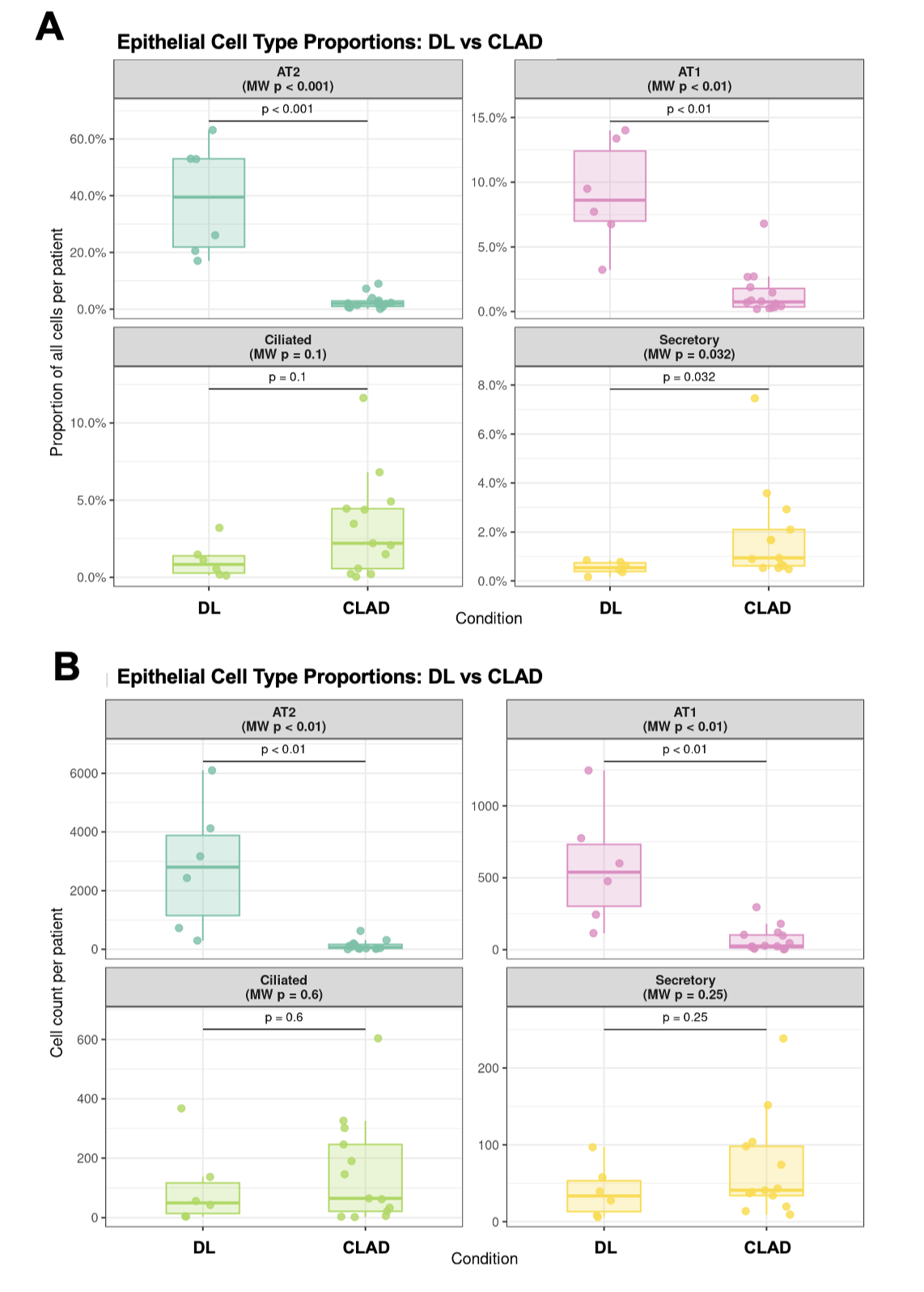

**Supplemental Figure 4. Per-patient epithelial cell composition in DL vs CLAD.** **(A)** Proportion of epithelial cell types per patient relative to all cells within each sample. **(B)** Counts of epithelial cell types per patient. Counts represent the number of cells per subtype within each sample from a given patient. AT2 cells represent the combined AT2 homeostatic, active, and transitional clusters. Each point represents an individual patient; boxplots summarize distributions (median, IQR, 1.5x whiskers). Statistical comparisons were performed using two-sided Mann-Whitney tests.

### **Supplemental MATERIALS AND METHODS**

*Single-cell RNA sequencing data analysis*

Samples subject to 3’ scRNA-seq were analyzed in R for Linux (v4.4.2) using Seurat (v5.2.1). Each sample was processed individually from filtered 10X matrices. Initial normalization, scaling, and Principal Component Analysis (PCA) were performed to support ambient RNA estimation and by correction with SoupX. Ribosomal protein genes were excluded prior to contamination estimation. The corrected counts were then used to generate new Seurat objects for each sample. Samples were filtered to retain cells with 200 to 5000 detected genes, 1000 to 30000 UMIs, and less than 15% mitochondrial reads. DoubletFinder was applied to each sample to retain only singlets. CLAD samples were merged into a single object, and DL samples, from a publicly available dataset *(1)*, were processed and merged in parallel. Batch correction was performed separately within each dataset using canonical correlation analysis-based integration. CLAD and DL objects were combined, followed by normalization, variable feature identification, data scaling, and PCA. Final batch correction was performed using reciprocal principal component analysis-based integration. Pre- and post-reperfusion public scRNA-seq data *(1)* were processed using the same workflow. For integration of publicly available IPF scRNA-seq datasets from Reyfman et al. *(2)*, the IPF object was generated using the same preprocessing workflow prior to Harmony-based integration with the CLAD/DL object.

Epithelial cells were isolated by subsetting clusters enriched for *EPCAM* expression and lacking the immune marker *PTPRC* from the fully integrated object. The epithelial subset was then re-processed independently – re-normalization, variable feature identification, scaling, and nearest-neighbor graph construction was performed using the pre-computed canonical correlation analysis reduction across 10 dimensions, followed by clustering at a resolution of 0.15 and generation of an epithelial-specific Uniform Manifold Approximation and Projection (UMAP) embedding. Epithelial subclusters were annotated based on marker gene expression.

To identify biologically relevant pathways, we performed over-representation analysis using the Hallmark gene set collection from the Molecular Signatures Database (MSigDB) via the *msigdbr* and *clusterProfiler* packages in R. Differentially expressed genes were first identified using the Wilcoxon rank-sum test implemented in Seurat's *FindMarkers* function. Significant genes were considered upon a minimum cell expression fraction of 25% and a log_2_ fold-change threshold of 0.25. Hallmark gene sets were assessed using a hypergeometric test with Benjamini-Hochberg multiple testing correction. Enriched terms were visualized as dot plots displaying gene ratio, gene count, and adjusted *p*-value for the top 20 significant pathways.

We performed RNA velocity analysis using *scVelo (3)*. Spliced and unspliced RNA count matrices were first extracted and matched to the annotated epithelial cell subset via barcode harmonization in R, using *Seurat*, *SingleCellExperiment*, and *zellkonverter*. Cell metadata and the UMAP embedding were retained. The finalized object was exported to H5AD format for downstream analysis in Python (v3.12.7). Counts were normalized and filtered retaining the top 2,000 highly variable genes with a minimum of 20 shared counts, after which PCA (30 components) and nearest-neighbor graph construction (30 neighbors) were performed using *scanpy*. Velocity streamlines were projected onto the existing UMAP embedding and visualized by cluster identity.

*Minor alloantigen mismatched mouse model of chronic lung allograft rejection*

Male C57BL/6 (B6, H-2^b^) recipient mice and C57BL/10 (B10, H-2^b^) donor mice (Jackson Laboratories, Bar Harbor, ME) were used in an OLT model with or without antecedent IRI as we previously described *(4)*. Briefly, donor B10 lungs were perfused with 4 °C LPD organ preservation solution and preserved at 4 °C for 6 hours (cold ischemic time, CIT). Warm ischemic time (WIT) consisted of a 45-minute incubation at 37 °C and 15 minutes for vascular and bronchial anastomosis (1-hour total WIT) prior to reperfusion. Recipients received 125 µL of EXP or control solution by intraperitoneal injection four times weekly beginning immediately after OLT and continuing until necropsy at 28-31 days post-transplant. At necropsy, the left (OLT) lung was divided into thirds.

One third of the left lung and the whole right (non-OLT) lung were seperately processed into single-cell suspensions for flow cytometry as described for the murine hilar clamp model of IRI. A second third was formalin-fixed for 24 hours, transferred to 70% ethanol, and paraffin-embedded for histologic assessment of graft injury as previously described *(4–8)*. Representative sections were stained with hematoxylin and eosin (H&E) and Masson trichrome (MT) by the UHN Pathology Research Program. The remaining third was snap-frozen and stored at -80 °C. For protein extraction, weighed frozen tissue was homogenized in 0.5 mL cold Mammalian Protein Extraction Reagent (Thermo Fisher) containing 1x HALT Protease and Phosphatase Inhibitor Cocktail and EDTA, using chilled stainless-steel beads in a TissueLyser II (Qiagen) for two 4-minute cycles at 20 Hz. Homogenates were held on ice for 20 minutes, cleared by centrifugation at 12,000 x g for 10 minutes at 4 °C, and the supernatant was stored at -80 °C. Murine SP-D was measured by ELISA (R&D Systems, DY6839-05) in cleared homogenate diluted 1:10 and normalized to total protein concentration by bicinchoninic acid assay.

*Statistical analysis*

Between-group differences in autoantibody reactivities on antigen microarrays were identified using SAM as previously described *(9)*, and isotype-specific average reactivities in BAL and serum were compared by two-way analysis of variance (ANOVA) with Šídák correction. Features shared between compartments were entered into greedy backward elimination with 5-fold cross-validation; within each fold, antibody-specific thresholds were set by Youden's J statistic and applied to held-out samples. Binarized features were summed into a composite score using our autoantibody scoring method as previously described *(9)*, and classifier performance was assessed by pooled out-of-fold ROC AUC.

Other two-group comparisons were assessed by Wilcoxon rank-sum tests. Kruskal-Wallis tests were used where more than two groups were compared. The latter additionally included Dunn's post hoc testing. Longitudinal biomarker data were summarized as mean ± SEM at discrete timepoints, without modeling longitudinal dependence; in human studies, samples obtained after the onset of CLAD were excluded and subjects were stratified by future CLAD status.

Analyses were performed in R for Mac (v4.2.1) or GraphPad Prism (v9.4.1). In R, rstatix was used for human longitudinal antibody nonparametric comparisons, pROC for ROC analysis and Youden’s J thresholding, and Seurat (R for Linux v4.4.2) for single-cell differential expression. Base R functions were used for Wilcoxon and Kruskal-Wallis tests, simple linear regression, and logistic regression for longitudinal antibody data, scRNA-seq, and primary disease antibody comparisons. Other analyses performed in GraphPad Prism included Kruskal-Wallis tests, unpaired two-tailed Student’s t-tests, two-way ANOVA, and simple linear regression.

References

1. A. Wong, A. Duong, G. Wilson, J. Yeung, S. MacParland, H. Han, M. Cypel, S. Keshavjee, M. Liu, Ischemia-reperfusion responses in human lung transplants at the single-cell resolution. *Am. J. Transplant.* **24**, 2199–2211 (2024).

2. P. A. Reyfman, J. M. Walter, N. Joshi, K. R. Anekalla, A. C. McQuattie-Pimentel, S. Chiu, R. Fernandez, M. Akbarpour, C.-I. Chen, Z. Ren, R. Verma, H. Abdala-Valencia, K. Nam, M. Chi, S. Han, F. J. Gonzalez-Gonzalez, S. Soberanes, S. Watanabe, K. J. N. Williams, A. S. Flozak, T. T. Nicholson, V. K. Morgan, D. R. Winter, M. Hinchcliff, C. L. Hrusch, R. D. Guzy, C. A. Bonham, A. I. Sperling, R. Bag, R. B. Hamanaka, G. M. Mutlu, A. V. Yeldandi, S. A. Marshall, A. Shilatifard, L. A. N. Amaral, H. Perlman, J. I. Sznajder, A. C. Argento, C. T. Gillespie, J. Dematte, M. Jain, B. D. Singer, K. M. Ridge, A. P. Lam, A. Bharat, S. M. Bhorade, C. J. Gottardi, G. R. S. Budinger, A. V. Misharin, Single-Cell Transcriptomic Analysis of Human Lung Provides Insights into the Pathobiology of Pulmonary Fibrosis. *Am. J. Respir. Crit. Care Med.* **199**, 1517–1536 (2019).

3. V. Bergen, M. Lange, S. Peidli, F. A. Wolf, F. J. Theis, Generalizing RNA velocity to transient cell states through dynamical modeling. *Nat. Biotechnol.* **38**, 1408–1414 (2020).

4. T. Watanabe, T. Martinu, A. Chruscinski, K. Boonstra, B. Joe, M. Horie, Z. Guan, K. F. Bei, D. M. Hwang, M. Liu, S. Keshavjee, S. C. Juvet, A B cell-dependent pathway drives chronic lung allograft rejection after ischemia-reperfusion injury in mice. *Am. J. Transplant. Off. J. Am. Soc. Transplant. Am. Soc. Transpl. Surg.* **19**, 3377–3389 (2019).

5. T. Watanabe, S. C. Juvet, G. Berra, J. Havlin, W. Zhong, K. Boonstra, T. Daigneault, M. Horie, C. Konoeda, G. Teskey, Z. Guan, D. M. Hwang, M. Liu, S. Keshavjee, T. Martinu, Donor IL-17 receptor A regulates LPS-potentiated acute and chronic murine lung allograft rejection. *JCI Insight* **8**, e158002.

6. S. Moshkelgosha, L. Levy, S. Safavi, S. Karunagaran, G. Wilson, B. Renaud-Picard, G. Madu, R. Ramchandani, J. Oliver, T. Watanabe, K. F. Bei, B. Joe, Q. Li, E. Huszti, M. Cheung, D. Hedley, J. Yeung, S. Keshavjee, T. Martinu, S. Juvet, Emergence of a senescent and inflammatory pulmonary CD4+ T cell population prior to lung allograft failure. *Sci. Adv.* **11**, eadp9052 (2025).

7. T. Martinu, H. Oishi, S. C. Juvet, M. Cypel, M. Liu, G. J. Berry, D. M. Hwang, S. Keshavjee, Spectrum of chronic lung allograft pathology in a mouse minor-mismatched orthotopic lung transplant model. *Am. J. Transplant. Off. J. Am. Soc. Transplant. Am. Soc. Transpl. Surg.* **19**, 247–258 (2019).

8. T. Watanabe, C. Lam, J. Oliver, H. Oishi, G. Teskey, S. Beber, K. Boonstra, J. Mauricio Umaña, H. Buhari, B. Joe, Z. Guan, M. Horie, S. Keshavjee, T. Martinu, S. C. Juvet, Donor Batf3 inhibits murine lung allograft rejection and airway fibrosis. *Mucosal Immunol.* **16**, 104–120 (2023).

9. D. Vosoughi, A. Ulahannan, Q. Li, E. Huszti, A. Chruscinski, D. Birriel, G. Madu, G. Teskey, M. Aversa, T. Martinu, S. Juvet, Humoral immunity to lung antigens early post-transplant confers risk for chronic lung allograft dysfunction. *J. Heart Lung Transplant.* **0** (2025), doi:10.1016/j.healun.2025.02.1577.
